# Linking candidate causal autoimmune variants to T cell networks using genetic and epigenetic screens in primary human T cells

**DOI:** 10.1101/2024.10.07.617092

**Authors:** Ching-Huang Ho, Maxwell A. Dippel, Meghan S. McQuade, Arpit Mishra, Stephan Pribitzer, LeAnn P. Nguyen, Samantha Hardy, Harshpreet Chandok, Florence Chardon, Troy A. McDiarmid, Hannah A. DeBerg, Jane H. Buckner, Jay Shendure, Carl G. de Boer, Michael H. Guo, Ryan Tewhey, John P. Ray

**Affiliations:** Benaroya Research Institute, Center for Systems Immunology; Seattle, 98101, USA; Benaroya Research Institute, Center for Translational Immunology; Seattle, 98101, USA; University of Washington, Molecular and Cellular Biology program; Seattle, WA, USA; The Jackson Laboratory; Bar Harbor, ME, USA; University of Washington, Department of Genome Sciences, Seattle, WA, USA; Seattle Hub for Synthetic Biology, Seattle, WA, USA; Howard Hughes Medical Institute, Seattle, WA, USA; University of British Columbia, School of Biomedical Engineering, Vancouver, BC, Canada; University of Pennsylvania, Department of Neurology, Philadelphia, PA, USA; University of Washington, Department of Immunology, Seattle, WA, USA

## Abstract

Genetic variants associated with autoimmune diseases are highly enriched within putative *cis*-regulatory regions of CD4^+^ T cells, suggesting that they alter disease risk via changes in gene regulation. However, very few genetic variants have been shown to affect T cell gene expression or function. We tested >18,000 autoimmune disease-associated variants for allele-specific expression using massively parallel reporter assays in primary human CD4^+^ T cells. The 545 expression-modulating variants (emVars) identified greatly enrich for likely causal variants. We provide evidence that many emVars are mediated by common upstream regulatory conduits, and that putative target genes of primary T cell emVars are highly enriched within a lymphocyte activation network. Using bulk and single-cell CRISPR-interference screens, we confirm that emVar-containing T cell *cis*-regulatory elements modulate both known and novel target genes that regulate T cell proliferation, providing plausible mechanisms by which these variants alter autoimmune disease risk.

## Main Text

CD4^+^ T cells play an essential role in immunity and autoimmune disease^1–4^, but how common genetic variants affect human CD4^+^ T cell function and disease pathology remain mostly unknown^5,6^. Genome-wide association studies (GWAS) have identified tens of thousands of genetic variants associated with autoimmune disease, but the vast majority are in non-coding regions and in tight linkage disequilibrium with many other variants^5,7,8^. Hence, >99% of causal variants that drive complex traits such as autoimmune diseases have yet to be determined^9^. Identifying causal variants and the cells in which they operate would aid in defining disease-relevant target genes, pathways, and cell types, and inform the development of more effective treatments^10^.

Complex trait-associated variants are enriched within chromatin that is accessible and/or actively modulating gene expression (i.e., within regions of H3K27ac deposited chromatin), which we define here as putative *cis*-regulatory elements (CREs). CREs differ between cell types^11^, and autoimmune GWAS variants enrich most highly within the CREs of CD4^+^ T cells^5,12–14^. Recent studies have shown that massively parallel reporter assays (MPRAs) can identify variants that alter *cis*-regulatory activity via changes in reporter gene expression^15–17^. Combining MPRAs with readouts of T cell-accessible chromatin enriched for causal variants up to 58-fold according to statistical fine-mapping^18^. Once putative causal variants are identified, a further challenge is to determine how they affect gene expression networks and cellular function. Online databases of functional genomic data such as Open Targets Genetics have been helpful for prioritizing genes that are likely targets of variants, but these databases lack perturbational data that directly link variants and CREs to gene expression^19^. To address this problem, recent studies have utilized bulk and single-cell CRISPR-interference (CRISPRi) screens to link variant CREs to the genes that they regulate^20–22^.

While the above experiments support the likely *cis*-regulatory role of GWAS variants in T cells and other blood cells, both MPRA and CRISPRi screens have rarely been applied within the context of primary cells. Instead, these experiments tend to employ immortalized cell lines such as a T cell leukemia cell line (Jurkat cells) or an erythroleukemia cell line (K562 cells). These cell lines may not accurately recapitulate the transcriptional regulation of primary cells or their associated phenotypes^23^. Testing these assays in primary cells containing transcriptional signatures and phenotypes that more closely reflect cells that play a role in disease would likely aid in highlighting the most relevant functional consequences of risk variants.

Here, we assayed variant function and connect variants to gene expression changes and cellular phenotypes within primary human CD4^+^ T cells. We tested >18,000 autoimmune variants for their ability to modulate CRE activity with MPRAs and found expression modulating variants (emVars) that act in primary T cells tend to be driven by inflammatory transcription factors and are largely distinct from those found in the Jurkat cancer cell line. We found primary T cell emVars enrich highly for causal variants, and that these variants are enriched within loci containing genes that act within a T cell activation gene network among other networks related to transcription and translation. Using single-cell and proliferation-based CRISPRi screens in primary T cells, we found that emVar CREs modulate genes within the T cell networks that control lymphocyte activation and mRNA processing, thus linking risk variants to T cell expression and function.

## Autoimmune variant MPRAs in primary human CD4^+^ T cells enrich for causal variants

We assessed the regulatory effects of genetic variants associated with multiple sclerosis (MS), type 1 diabetes (T1D), psoriasis, rheumatoid arthritis (RA), and inflammatory bowel disease (IBD)^24–28^ in activated primary human CD4^+^ T cells using MPRA (Fig. 1a). We used a library previously described for a study in Jurkat T cells totaling 578 index single nucleotide polymorphisms (SNPs) and 18,312 total variants in tight linkage disequilibrium (r^2^>0.8) in the European subset of the 1000 Genomes Project cohort^18^.

**Fig. 1.**
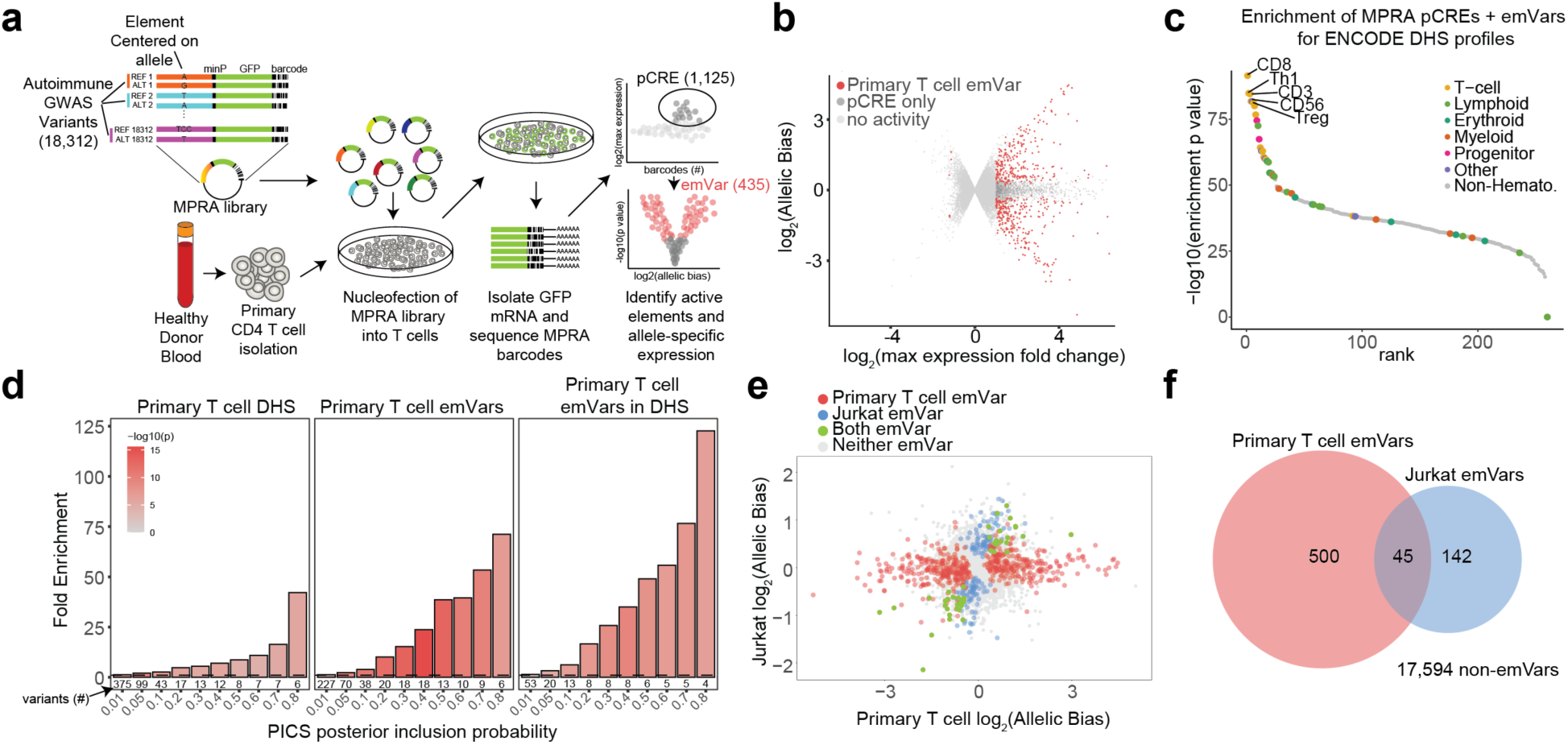
Autoimmune-associated emVars enrich for autoimmune disease-causal variants. **a**, Primary T cell MPRA workflow. **b,** Volcano plot. The log_2_ expression of the highest expressing allele is on the x axis and the log_2_ of the activity of allele1/allele2 is on the y axis. **c,** Enrichment of pCRE and emVar elements for the accessible chromatin profiles of all ENCODE cell types. The –log_10_ *P* value for the enrichment of pCRE and emVar elements is on the y axis and the rank according to this *P* value is on the x axis. Cell lineages are depicted according to the colors in the legend. **d,** Bar plot showing the enrichment of DHS elements, emVars identified in primary T cells, and emVars in DHS elements for PICS statistically fine-mapped variants using probability thresholds indicated on the x axis. Bar plot shade indicates –log_10_ *P*-value enrichment. Numbers below bars indicate the number of emVars that are statistically fine-mapped at a given PICS probability. Enrichment was calculated as a risk ratio, with *P*-values determined through a two-sided Fisher’s exact test. **e,** Scatterplot comparing element allelic skew between Jurkat and primary T cell MPRA libraries. The log_2_ allelic bias levels of MPRA elements tested in primary T cells are plotted on the x-axis and in Jurkat on the y-axis. **f,** Venn diagram depicting called emVars between primary T cell and Jurkat MPRAs.

We observed high reproducibility of the primary T cell MPRA across independent donors (Pearson correlation > 0.93; Supplementary Fig. 1a-c). We identified MPRA elements that have CRE activity greater than the baseline activity of the minimal promoter (Supplementary Figs. 1d and 2a, b), identifying 1125 putative CREs (Supplementary Fig. 2c, d; Supplementary Tables 1 and 2; Methods). Of these putative CREs, we identified 545 expression-modulating variants (emVars) according to significant differences in allele-specific expression (Fig. 1b; Supplementary Table 1). We found primary T cell emVars in one third of tested loci (Supplementary Fig. 3a), with only one emVar on the haplotype in 53.6% of tested loci (Supplementary Fig. 3b). Primary T cell emVars and putative CREs were both enriched within active chromatin marks and other readouts of *cis*-regulatory activity, and were found preferentially in the accessible chromatin of primary T cells compared to other cell types (Fig. 1c; Supplementary Fig. 4; Supplementary Note; Supplementary Tables 3-5)^18,29^. Thus, primary T cell emVars and putative CREs are enriched within endogenous T cell regulatory elements.

To determine whether primary T cell emVars enrich for causal variants, we assessed emVar enrichment for causal variants according to both Probabilistic Identification of Causal SNPs (PICS) posterior inclusion probabilities (PIPs) for each trait locus^5,18^ and SuSIE fine-mapping data for UK Biobank traits (Fig. 1d; Supplementary Fig. 5; Supplementary Tables 6 and 7)^30^. In loci where at least one emVar is observed, emVars with high posterior probabilities were enriched upwards of 71-fold (PICS) and 50-fold (UKBB) compared to all other tested variants (Fig. 1d, center; Fig. 5c, center; Supplementary Tables 8 and 9). This enrichment was further increased when considering only variants in T cell DNase Hypersensitivity (DHS) sites with 122-(PICS) to 200-fold (UKBB) enrichment for high posterior probability variants (Fig. 1d, right; Supplementary Fig. 5c, right; Supplementary Table 8 and 9). Overall, we found that our primary T cell MPRAs had a sensitivity of 27% and a specificity of 88% for identifying variants in fine-mapped 95% credible sets in risk loci. Thus, primary T cell emVars in T cell accessible chromatin enrich highly for fine-mapped causal variants.

## Transcription factor usage varies by cell type and disease

Although both primary T cell and Jurkat emVars enrich highly for causal variants (Fig. 1d; Supplementary Fig. 5)^18^, only 45 emVars overlapped between datasets, albeit this was significantly more than expected by chance (hypergeometric *P* = 5.4e-29). Primary T cell and Jurkat MPRAs differed largely in activity and allelic bias in reporter expression (Fig. 1e,f), thus we hypothesized that differences in signaling and activation between cell types may alter the transcription factor (TF) programs operating in each cell type^23,31^. To determine whether there were differences in the TFs acting in T cells and Jurkat cells, we first identified TFs responsible for MPRA activity. We assessed how each variant is predicted to change TF binding using known catalogs of TF motifs and compared this to the allelic effect on reporter expression within MPRA (Methods). We identified TF motifs that, when disrupted by variants, reduce expression in the MPRA, suggesting that these are transcriptional activators, such as ATF1, and conversely, variant-disrupted motifs with a corresponding increase in expression, suggesting that these are transcriptional repressors, such as GFI1B, consistent with GFI1B’s known repressor role (Fig. 2)^32^. While many TF effects were shared between both T cells and Jurkat cells, variants that disrupted transcriptional activators associated with inflammation were specific to primary T cells, including NFKB1 (n=67 variants), STAT3 (n=71 variants), and JUN and FOSB (n=72 and 61 variants, respectively) (Fig. 2 and Supplementary Fig. 6a and b; Supplementary Tables 10-13). TF motifs whose perturbation by variants had concordant effects on MPRA expression in both cell types include ATF1, ETS factors ELK1 and 4 and ELF1 and 2, and GFI1B (Fig. 2 and Supplementary Fig. 6a and b; Supplementary Tables 10-13). Thus, while both Jurkat and primary T cell MPRAs enriched for causal variants, differences in emVar identification in each setting are likely driven by alternative cellular programs and orchestrated by TFs.

**Fig. 2.**
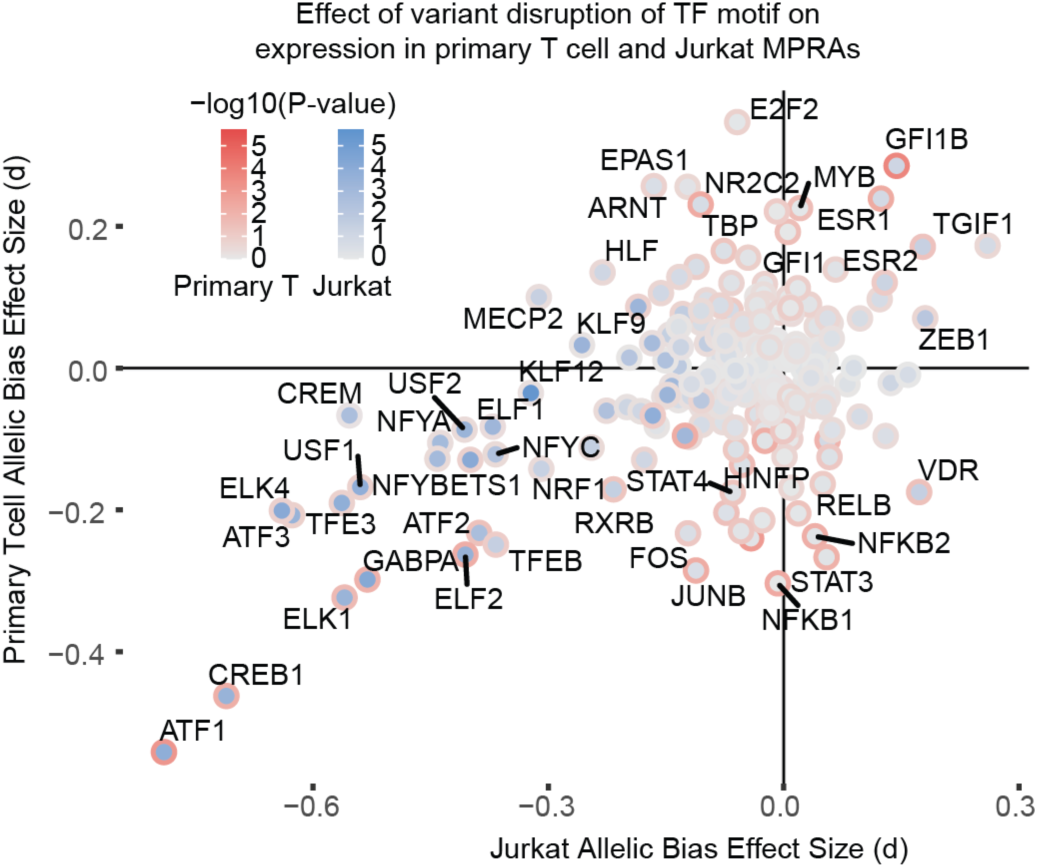
Differences in predicted transcription factor motif disruption between Jurkat and primary T cell MPRA libraries. Scatterplot comparing the effect of variants that are predicted to disrupt a TF motif and subsequent cumulative effect on MPRA expression between both primary T cell (red outline) and Jurkat (blue fill) libraries. The effect size is calculated using Cohen’s d for variant alleles predicted to disrupt a given TF motif and *P*-values are calculated using a *t* test comparing effect on expression of variants that disrupt a given motif versus all other variants.

To determine whether we can identify TFs that drive risk in a disease-specific manner, we grouped variants by disease and repeated the analysis. We observed several TFs with disease-specific enrichment, including the ZNF563 motif, whose disruption is highly activating at IBD (d = 0.38, *P* = 0.0002) and RA loci (d = 0.47, *P* = 0.12), but repressive at psoriasis loci (d = –0.84, *P* = 0.005), and disruption of the GATA3 motif to be highly activating at MS loci (d = 0.32, *P* = 0.001) but with an average of no effect in loci associated with other diseases (Supplementary Fig. 6c). Thus, we find T cell MPRAs to be sensitive to TF usage, providing direct evidence for the importance of performing MPRAs across multiple cellular contexts for defining causal variants, and we define TFs that may be more important at specific disease loci.

## emVars connect to T cell regulatory networks via multiple pathways

Motivated by the observation that the critical transcriptional regulators of T cell responses appear to mediate some primary T cell emVars, we hypothesized that emVars could increase disease risk by modulating the expression of genes that are critical for T cell responses. To test this, we compared putative target genes of emVars in primary T cell DHS sites that were identified in primary T cell versus Jurkat MPRAs. We identified 79 primary T cell and 31 Jurkat emVars in T cell DHS sites (10 emVars overlapping) within the Open Targets Genetics Variant to Gene (V2G) dataset^19^. Each emVar was associated with an average of 8-11 genes for a total of 671 (primary T cell) and 336 (Jurkat) emVar-linked genes that are expressed in T cells with a TPM>1 (230 genes shared between datasets; chi squared *P* < 2.2e-16; Supplementary Tables 14 and 15). We input these genes into STRING^33^ to define a primary T cell network of genes according to gene interaction experiments, co-expression, and text mining (Methods). The resulting primary T cell network was more highly connected than expected when compared to a background of all V2G genes linked to all MPRA-tested variants in T cell DHS sites (STRING PPI enrichment *P* < 1e-16; Fig. 3a; Supplementary Data 1). Overall, the primary T cell network was enriched for T cell activation according to EnrichR (false discovery rate (FDR) = 0.026, Panther module; Supplementary Table 16). We then defined clusters within the network and observed that the largest clusters were involved in lymphocyte activation, translation, transcriptional regulation, antigen processing, mRNA processing, and mRNA splicing (Fig. 3a and b; Supplementary Table 17).

**Fig. 3.**
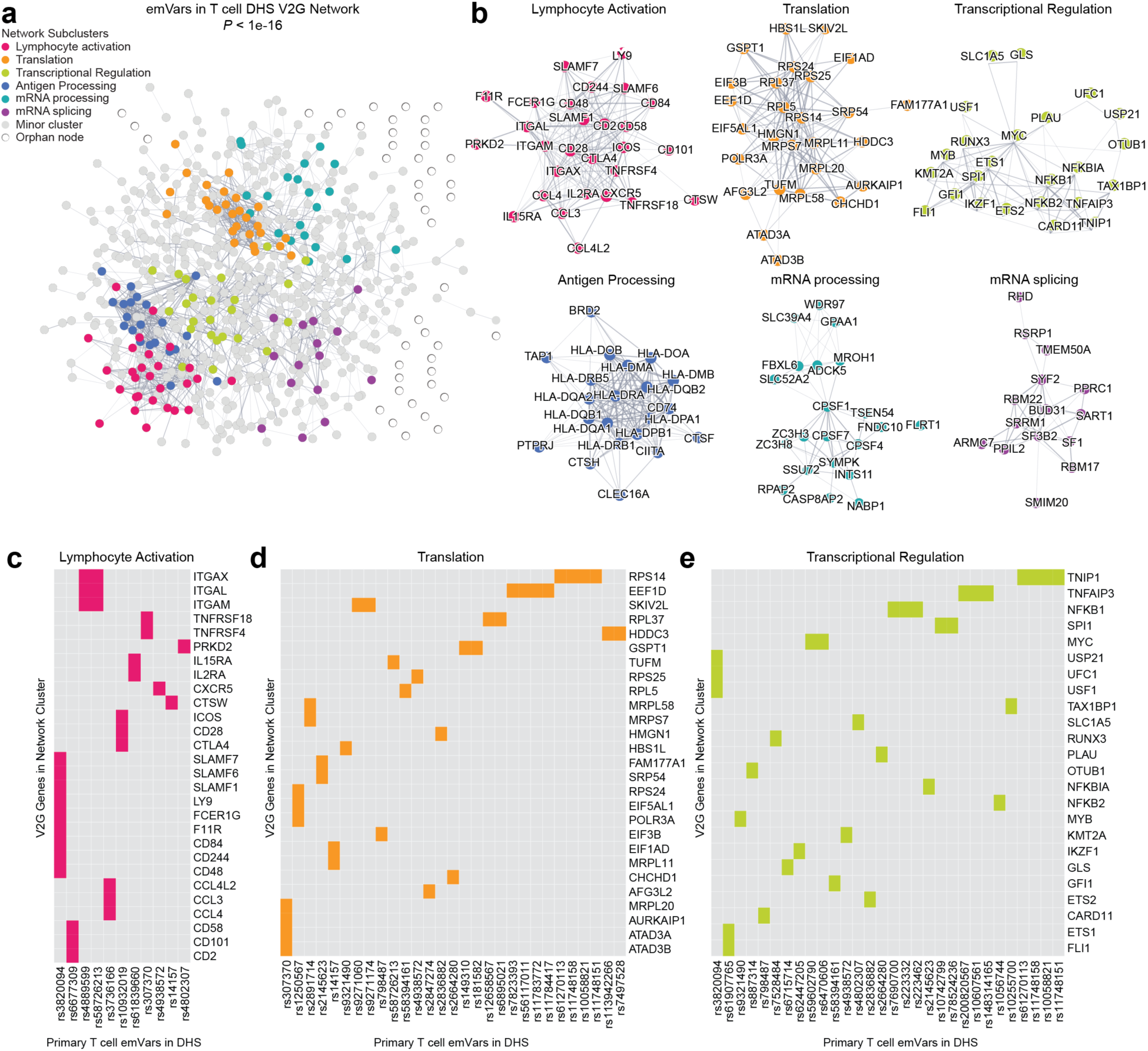
Network analysis of predicted target genes of emVars identifies significant enrichment of T cell activation cluster. **a**, STRING network showing V2G genes linked to 79 emVars in T cell DHS sites (nodes) and edges representing the strength of gene-gene interactions. **b,** The subclusters with the most genes from the larger network in (a) with labeled gene nodes. **c-e,** The lymphocyte activation (c), translation (d), and transcriptional regulation (e) clusters with each emVar on the x-axis and target gene on the y-axis. Fill color indicates that the gene is a V2G gene of the indicated emVar. *P*-value in (a) is calculated according to STRING protein-protein interaction enrichment given the expected versus observed number of edges within the network.

Within the lymphocyte activation cluster were known costimulatory genes expressed in T cells that regulate T cell activation including *CD28, CTLA4, ICOS, TNFRSF18* (GITR)*, TNFRSF4* (OX40), and *SLAM* family members (Fig. 3B, pink cluster). We also found that the transcriptional regulation cluster contained several members of the NF-κB signaling family *NFKB1, NFKB2, NFKBIA, TNFAIP3,* and *TNIP1* (Fig. 3B, yellow-green cluster). We compared these clusters to the network built on V2G genes linked to Jurkat emVars in T cell DHS sites finding that, while there was a cluster containing immune-relevant signaling molecules, there were no T cell costimulatory genes within this or any other cluster (Supplementary Fig. 7a and b, pink cluster; Supplementary Table 18). Furthermore, we found a transcriptional regulation cluster within the Jurkat network but this did not contain any members of the NFKB signaling axis (Supplementary Fig. 7a and b, yellow-green cluster; Supplementary Table 18). Thus, the putative target genes of emVars found in primary T cell libraries are more involved in T cell costimulation and NF-κB signaling than those of Jurkat emVars.

To link the emVars back to the putative genes they regulate within each network, we created an emVar by gene matrix for each cluster (Fig. 3c-e; Supplementary Fig. 7c; Supplementary Fig. 8a-c). Interestingly, while some emVars in DHS sites highlighted by both the primary T cell and Jurkat clusters were shared, such as rs58726213, which putatively regulates integrin genes *ITGAL, ITGAX,* and *ITGAM* (Fig. 3c; Supplementary Fig. 7c pink cluster), we also identified different emVars in each assay that putatively regulate these genes (rs4889599 – primary T cell, Fig. 3c; rs6565217 – Jurkat, Supplementary Fig. 7c pink cluster). Most emVar:gene interactions in the top 5 clusters for each cell type do not overlap (26/104 and 26/222 shared emVar:gene interactions in Jurkat and primary T cell networks, respectively), suggesting that different emVar:gene interactions drove each cluster. Collectively, emVars identified using primary T cell MPRAs occur more frequently in loci containing costimulatory and inflammatory genes compared to emVars identified in Jurkat cells.

## Single-cell CRISPRi screens reveal multiple pathways from variant to regulatory network

While V2G data provide putative gene targets of variants, we sought to connect variants directly to the genes they regulate using a single-cell CRISPR-interference (scCRISPRi) approach in primary T cells. We used guide RNAs (gRNAs) to target dCas9 tethered to an inactivating ZIM3-KRAB domain to variant CREs and assessed local effects on gene expression (within 1 Mb) with single cell RNA-sequencing^34^. We created two gRNA libraries to test 56 total emVars in T cell DHS sites. The first library targeted 20 emVars and 3 putative CREs, prioritizing emVars more likely to be causal variants by PIP. The second library targeted 50 T cell emVars in T cell DHS sites > 3,500 bp from transcription start sites (TSSs), as to avoid silencing of promoter regions. 13 emVars overlapped between both libraries (Fig. 4a; Supplementary Tables 19 and 20). We used SCEPTRE^35^ to connect CREs to local genes (<1Mb) based on differential gene expression in cells containing gRNAs targeting the CRE versus those containing non-target gRNAs.

**Fig. 4.**
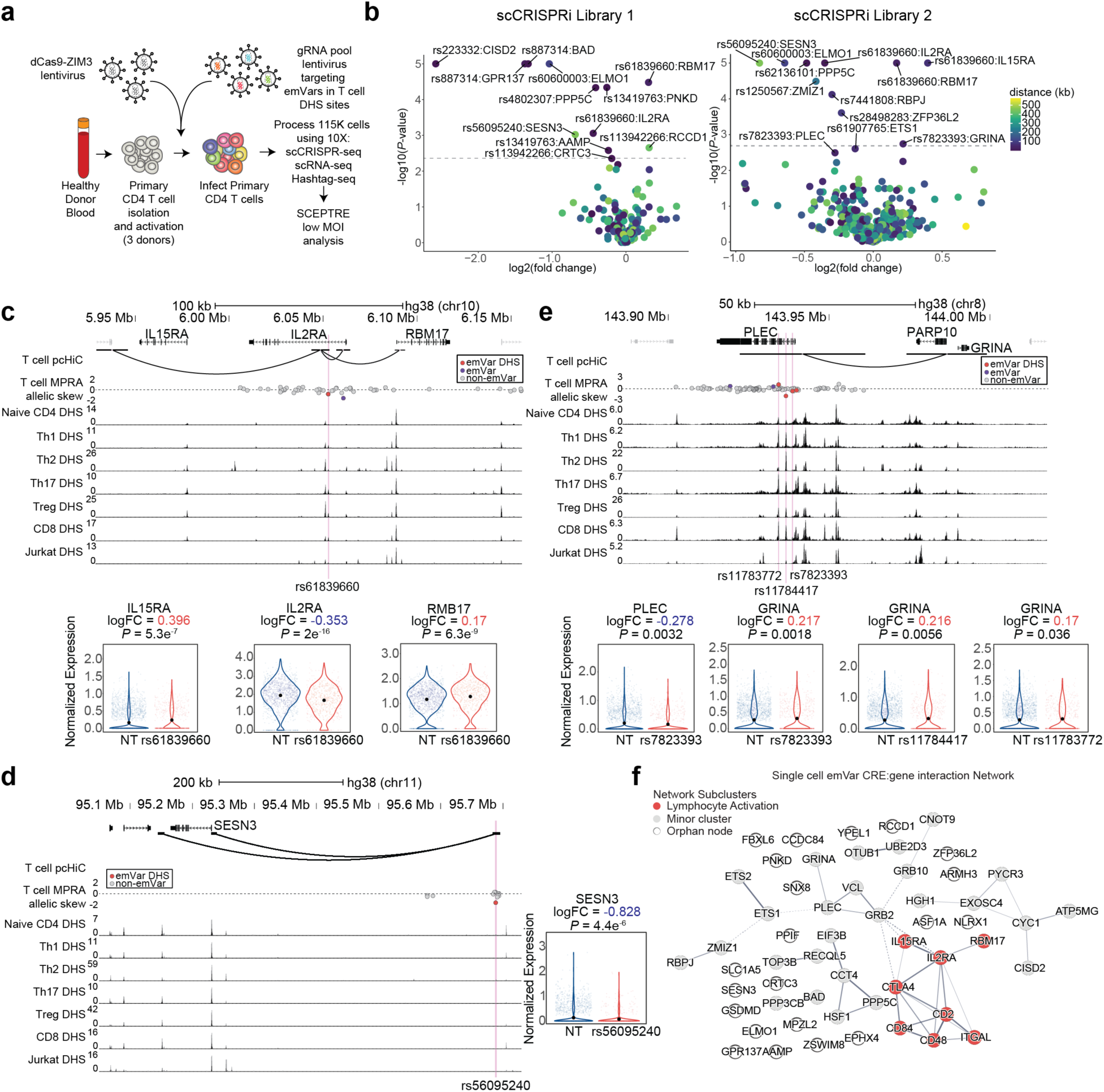
Single cell CRISPR-interference screens connect emVars to target genes. **a**, Workflow for scCRISPRi screens. **b,** Volcano plots depicting significantly differentially expressed genes when targeting a given emVar CRE. Log_2_ fold change is on the x-axis and –log_10_ *P* value for differential gene expression is on the y-axis. *P* values were calculated according to SCEPTRE. Dotted line indicates the empirical significance cutoff determined by SCEPTRE based on calibration with the non-target control gRNAs. **c-e,** Locus plot of the IL2RA (c), SESN3 (d), and PLEC loci (e). pcHiC loops from primary human T cells are depicted below genes in the locus plot. Disease-associated variants (dots) are red if they are emVars in T cell DHS sites, blue if they are emVars not within DHS sites, and gray if they are non-emVars. Accessible chromatin data from T cells are depicted as read pileups (peaks) on the locus track from various T cell types. The pink lines represent the location of emVars in DHS. Violin plots depict genes that are differentially expressed when targeting CRISPRi to the emVar using gRNAs compared to cells containing non-target gRNAs. **f,** SCEPTRE network of genes identified using scCRISPRi screens compared to all tested genes for 56 emVars in accessible chromatin. The red subcluster indicates the lymphocyte activation network. Black dots in violin plots in (c, d, and e) depict the mean of the distribution.

We found 13 of the 56 tested emVars to impact at least one gene in cis, with a total of 18 significant emVar CRE:gene interactions (Fig. 4b; Supplementary Fig. 9a, b; Supplementary Tables 21 and 22). Among them, we found rs61839660 (T1D and IBD PICS=0.98), an *IL2RA* intronic variant (9kb from TSS) previously associated with the timing of *IL2RA* expression in murine T cells^36^, was associated with a downregulation of *IL2RA* but also an upregulation of several nearby genes including *IL15RA*, a gene involved in homeostatic proliferation of memory T cells, and *RBM17*, involved in non-sense mediated decay (Fig. 4b and 4c)^37^. We also identified rs887314 (psoriasis PICS = 0.13) within the promoter of *BAD*, which we found to not only regulate *BAD*, a gene involved in T cell development and apoptosis^38^, but also *GPR137*, and *PYGM* expression (Fig. 4b; Supplementary Fig. 9c). Other notable hits include rs56095240 (MS PICS = 0.18) in an intergenic region 456 Kb from the *SESN3* TSS, which regulates *SESN3* expression (Fig. 4b, d), a gene involved in negative regulation of reactive oxygen species signaling ^39^, albeit its function has yet to be determined in T cells, and rs60600003 (MS PICS = 0.41), in intron 1 of *ELMO1*, 106 Kb from the *ELMO1* TSS, which had a substantial effect on ELMO1 expression, a gene involved in lymphocyte motility (Fig. 4b, left; Supplementary Fig. 9d)^40^. In addition, two other emVars, rs1250567 (MS PICS = 0.03) and rs7441808 (RA PICS = 0.007 and eosinophil counts UKBB PIP = 0.18) were significantly associated with *RBPJ* and *ZMIZ1*, respectively, both involved in WNT signaling in T cells (Fig. 4b, right)^41,42^, and rs61907765 (Psoriasis PICS = 0.43) was associated with *ETS1* expression, a TF involved in survival and activation of T cells and the development of natural regulatory T cells (Fig. 4b, right)^43^. We also identified genes that could play a role in disease biology with no previous evidence of contributing to T cell biology. For example, targeting three emVars in separate CREs in a large haplotype within *PLEC* with CRISPRi lead to an upregulation in *GRINA* expression, a glutamate receptor (Fig 4e); neither *PLEC* nor *GRINA* have an established role in T cells. Furthermore, between both screens, we identified two emVars that regulate *PPP5C* expression, a phosphatase that acts downstream of ERK signaling^44^, but that has no established role in T cell biology (Fig. 4b).

Finally, to assess if genes identified by our scCRISPRi screens were enriched in T cell related networks, we relaxed our calling threshold to include variant CRE:gene interactions of marginal significance (*P < 0.05)*. We were able to connect 41 of the 56 tested variants (71.9%) to 67 genes, of which 65 overlapped with V2G genes (Supplementary Tables 21 and 22). Interestingly, through creating a STRING network based on the hits versus all local genes that were tested in the single-cell screen, we again found that the T cell activation cluster was the most predominant cluster in the network (Methods; Fig. 4f; Supplementary Table 23), further supporting the importance of T cell activation programs in genetic risk for autoimmunity.

## Identification of variant CREs that regulate T cell proliferation

While many primary T cell emVar target genes were found within or connected to T cell activation networks, whether emVar CREs actually impact T cell proliferation remains unknown. To systematically link variant CREs with T cell activation and proliferation, we employed bulk CRISPRi screens in primary human T cells (Fig. 5a). Because these data could be broadly useful for classifying variant function, we also assessed ∼1000 additional autoimmune variant CREs (Supplementary Fig. 10a, b). We created a gRNA library targeting on average 14 gRNAs to each variant CRE (within 100 bp of each variant), 120 positive control gRNAs targeting known regulators of T cell proliferation and effector function, and 992 non-targeting control gRNAs (Fig. 5a; Supplementary Table 24). We activated primary human T cells and delivered dCas9-ZIM3 and the gRNA library using lentivirus infection. We harvested half of the transduced cells directly post-puromycin treatment (day 2) and the remainder of the cells after 21 days of proliferation (day 21). Integrated gRNA spacer sequences were sequenced and found to be enriched or depleted in day 21 compared to day 2 using MAGeCK^45^.

**Fig. 5.**
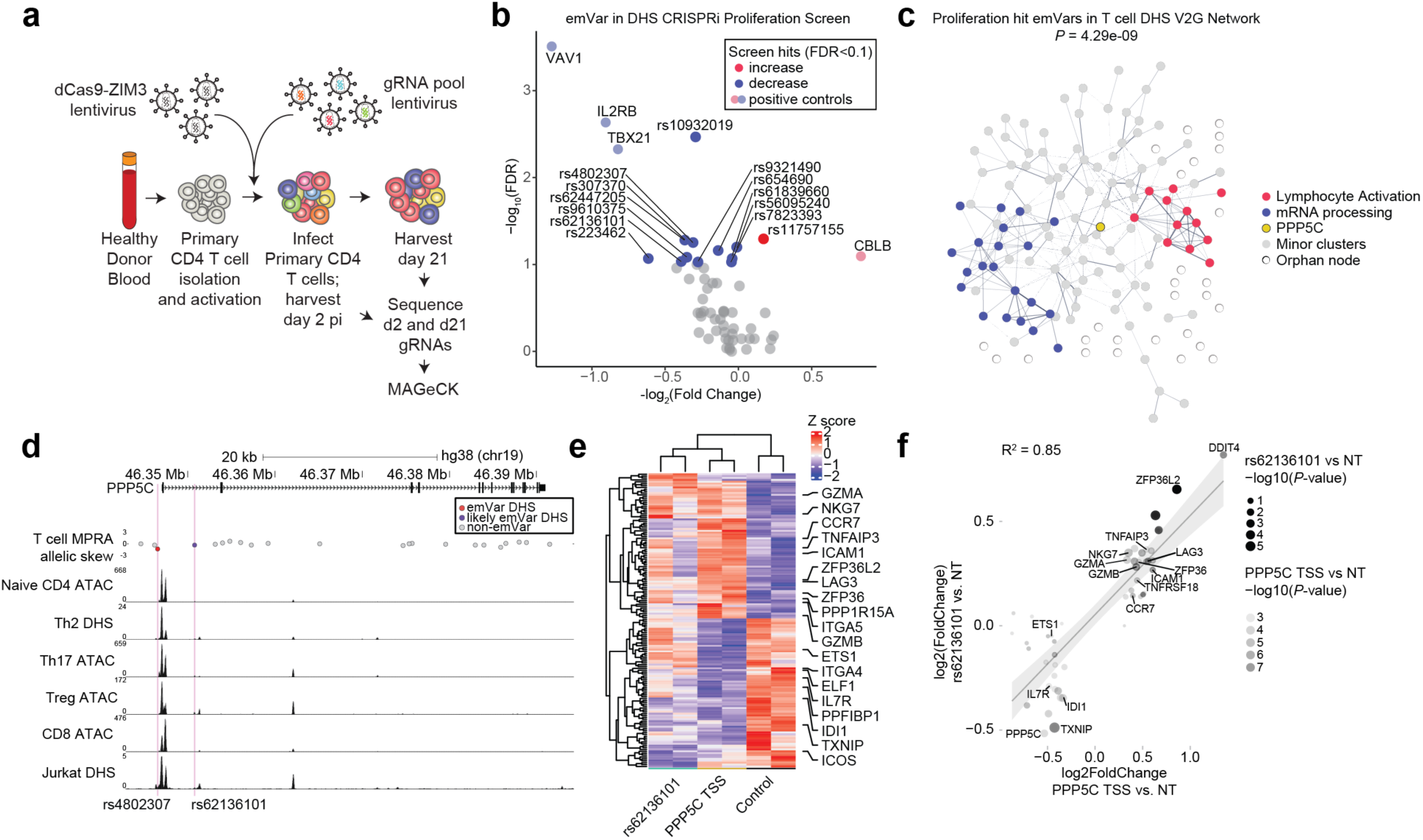
Proliferation screens identify emVars that modulate T cell proliferation. **a**, Proliferation screen experimental workflow. **b,** Volcano plot of significant positive control genes and variant CREs (blue and red) and non-significant targets (gray), with the log_2_ fold-change on the x-axis and the –log_10_ false discovery rate on the y-axis. **c,** STRING network based on 13 emVars in DHS sites that are CRISPRi proliferation hits. The lymphocyte activation and mRNA processing clusters and the PPP5C gene are highlighted in color. **d,** Locus plot depicting the PPP5C locus. pcHiC loops from primary human T cells are depicted below genes in the locus plot. Disease-associated variants (dots) are depicted: rs4802307 (red), an emVar in a T cell DHS site, rs62136101(blue), a likely emVar in a T cell DHS site, and non-emVars (gray). Accessible chromatin data from T cells are depicted as read pileups (peaks) on the locus track from various T cell types. The pink lines represent the location of emVars in DHS sites. **e,** Heatmap of differentially expressed genes when targeting the PPP5C TSS or rs62136101 with CRISPRi using gRNAs compared to cells containing a non-target gRNA. **f,** Scatterplot depicting the correlation between differentially expressed genes when targeting rs62136101 vs. NT (y axis) the PPP5C TSS versus non-target (NT; x axis). *P*-values in (f) are determined using a Wald test with DESeq2. Shaded region in (f) is the 95% confidence interval.

Through analyzing the effect of targeting all ∼1000 autoimmune GWAS variants in T cell DHS sites, we identified known positive controls, including VAV1 and IL2RB as positive regulators of T cell proliferation, and, CBLB, a known negative regulator (Supplementary Fig. 10c; Supplementary Table 25)^46,47^. We identified 21 additional variant CREs that were significantly associated with T cell proliferation (padj<0.1; Supplementary Fig. 10c; Supplementary Table 25). Among the hits with the strongest positive effect were variant CREs that contact the MYC promoter according to promoter capture HiC in T cells, such as rs10098999 (RA PICS = 0.0014) and rs10113762 (RA PICS = 0.0145), both of which are 800 kb downstream of MYC (Supplementary Fig. 10d) ^48^. To determine whether our CRISPRi proliferation screen hits were enriched for causal variants, we again used a risk ratio to determine enrichment for PICS fine-mapped variants, finding no enrichment for causal variants (PICS > 0.1: risk ratio enrichment = 0.52, Fisher’s exact test *P* = 0.57), even when relaxing the FDR threshold to 0.2 (risk ratio enrichment = 0.59; Fisher’s exact test *P =* 0.33). Thus, we did not find that our CRISPRi proliferation screens enriched for causal variants and therefore chose to focus our analysis on variants that are primary T cell emVars, which we have already found to be enriched for causal variants (Fig. 1d).

Focusing our analysis on the 56 emVar CREs that we analyzed in our scCRISPRi screens, we analyzed gRNAs that were enriched or depleted at day 21 compared to day 2 (Fig 5b; Supplementary Table 26) identifying 12 emVar CREs that reduce proliferation when targeted with CRISPRi, and 1 emVar CRE that promotes proliferation. Interestingly the most significant emVar CRE hit that reduced T cell proliferation when targeted, rs10932019 (RA PICS 0.007), is 50kb downstream from *CD28*, required for T cell activation. Other emVar CREs of note that reduced T cell proliferation when targeted were rs307370 (IBD PICS = 0.018), in the *TNFRSF4* (encoding OX40) locus, rs61839660 (T1D PICS = 0.98) in the *IL2RA* locus, and rs9610375 (MS PICS = 0.0028) in the *MAPK1* locus; each of these genes has previously been shown to be a positive regulator of T cell proliferation^49–51^. Conversely, the emVar CRE that increased T cell proliferation when targeted, rs11757155 (IBD PICS = 0.017) is in an intron of *BACH2*, a transcription factor involved in suppressing T cell activation and effector T cell differentiation^52^. Given that many of our emVar CRE hits appeared in the T cell signaling and proliferation networks highlighted in the MPRA STRING network (Fig. 3), we next sought to determine whether the V2G genes of emVar CRE proliferation hits were enriched for these network clusters compared to a background of V2G genes linked to all emVar CREs tested in the proliferation screen (Supplementary Table 27). We again found the top represented clusters were those pertaining to lymphocyte activation and signaling along with mRNA processing (Fig. 5c; Supplementary Fig. 11a, b; Supplementary Table 28). Our scCRISPRi screens found that 5 of the 13 proliferation hits also had a significant effect on the expression of local genes, including rs61839660 with IL2RA, IL15RA, and RBM17, rs7823393 with GRINA, rs56095240 with SESN3, and both rs4802307 and rs62136101 with PPP5C (Fig. 4b). Thus, we found that 13 of the 56 tested emVar CREs (23%) had a significant effect on T cell proliferation, linking these putatively causal variants to T cell function, and highlighting putative target genes that enrich for T cell activation pathways.

## PPP5C is a novel regulator of T cell proliferation

Two emVar CREs that reduced T cell proliferation when targeted, rs4802307 (emVar) and rs62136101 (an emVar with allelic skew in 6 out of 7 donors; Supplementary Fig. 11c), are in the PPP5C locus on the same haplotype. The rs4802307-A (IBD lead variant) and rs62136101-T (r^2^=0.96 to rs4802307 in Europeans) alleles are associated with reduced PPP5C expression in T cell expression quantitative trait locus (eQTL) data and are protective from IBD (IBD PICS = 0.1 and 0.06, respectively) (Fig. 5d) ^28^. From our single cell screens, we found that *PPP5C* is the only significantly differentially expressed gene locally, suggesting this is the target gene. PPP5C promotes RAF-MEK-ERK signaling and cancer cell proliferation^44,53^, but its function in T cells has not been assessed. Because we find PPP5C affects T cell proliferation but is not directly within the main T cell activation network, we hypothesized that PPP5C could be a distal regulatory node to the T cell activation cluster. To assess this, we determine global programs that are differentially regulated upon targeting CRISPRi to the rs62136101 CRE and the *PPP5C* TSS in T cells (Fig. 5e; Supplementary Tables 29 and 30), finding an upregulation of genes associated with T cell metabolism and function. These including *ZFP36L2*, which has been shown to regulate IFNg production and homeostatic and autoreactive T cell responses^54,55^, *DDIT4*, which suppresses mTOR function^56^, *LAG3* which suppresses T cell activation^57^, and granzymes *GZMA* and *GZMB* which are used by cytotoxic and regulatory T cells to promote or suppress immune responses, respectively^58,59^. While targeting CRISPRi to rs62136101 had a more subtle effect than the TSS, both led to consistent differentially expressed genes involved in T cell biology (Fig. 5f). Thus, PPP5C is a previously unappreciated component of T cell signaling, which IBD protective alleles downregulate leading to tuning of T cell metabolic and effector programs.

## Discussion

Identifying variants that underlie complex traits and defining their effects on disease-relevant cell types continues to be a longstanding challenge. Through dissecting autoimmune disease-associated variant function using genomic screens and features in human primary CD4 T cells, we identified likely causal variants with high precision, narrowed their potential target genes, and gained insights into pathways that causal variants regulate. Indeed, we show many examples of candidate causal variants in regulating genes involved with T cell signal transduction, translation, and RNA processing, suggesting that these are key pathways within T cells that contribute to autoimmune disease genetic risk. These data suggest that performing assays across other relevant primary cell types will unveil other causal variants and pathways at play within autoimmune diseases, helping to unlock mechanisms and potential therapeutic targets.

Identification of causal variants that alter CRE activity requires testing their effects in relevant cell types. Primary T cells underlie the pathogenesis of many autoimmune diseases as shown, for example, by effective therapeutics targeting CD3 that treat T1D^60^ and targeting the IL23 pathway in IBD and psoriasis^61,62^, and by the preferential enrichment of autoimmune disease-associated genetic variants within T cell-accessible chromatin^7,13,14^. However, primary T cells have been notoriously difficult to genetically engineer and perturb until recent advances^47,63^. The Jurkat cell line has served as a tractable model for primary T cells for decades, though they differ in a number of ways, with mutations in key tumor suppressors such as P53, the PI3K pathways, large structural variants, and heightened expression of MYC^23,64^. Through testing the same MPRA library in both primary T and Jurkat cells, we find that both settings identify causal variants, but the emVars identified differ largely^18^. While these data suggest that both contexts are relevant for defining causal variants, we find that their differences are likely controlled by a unique suite of TFs, with inflammatory TFs playing a predominant role in primary T cells. Interestingly, we also find that the cellular context in which MPRA is conducted seems to be relevant to the specific loci that contain emVars, as demonstrated by the enrichment of primary T cell emVar candidate target genes for T cell co-stimulatory molecules and NF-κB family members. These data suggest that testing this MPRA library in more disease-relevant cellular contexts, including other T cell subsets, B cells, myeloid cells, and non-immune cells, will identify additional causal variants in other loci orchestrated by unique upstream regulators.

Once causal variants are identified, there remains a key challenge in defining their target genes and pathways. Others have begun to do this using bulk and scCRISPRi screens in cell lines, particularly within the K562 cell line^20,22^. Our scCRISPRi screens in primary T cells identify emVar target genes that operate within the same T lymphocyte activation network that is highlighted within our V2G-based network analysis. This is in spite of the fact that single cell screens are still less likely to pick up smaller effect changes in expression. Additional target genes might be identified through increasing the number and efficacy of gRNAs and increasing the number of cells assayed. However, the effect size of a given enhancer on a gene and the specific context in which an enhancer functions on a gene will also affect whether a target gene is identified in these screens^65^. In some cases, we find that emVars regulate more than one gene. Thus, disentangling causal variant effects will likely be more complex than focusing on singular target genes within each locus, which has been the focus of genetic knockout studies for many years.

While we expected causal variants to have larger effects on T cell function, our genome-wide CRISPRi screen of ∼1000 variant CREs for their effects on T cell proliferation did not seem to enrich for causal variants. Given these data, we suspect that variants capable of imposing large effects on regulatory regions important for directly regulating disease processes are more likely to be rarer or eliminated in populations through purifying selection. Inversely, common disease variants may be more likely to reside in low-impact enhancers of key disease genes or higher-impact enhancers of non-critical genes with modest effects on cellular function. These data are in line with the recent observations showing systematic differences between eQTLs and GWAS loci where disease variants that have a high impact on a trait tend to have lower effects on local gene expression^66^. Through integrating our MPRA, scCRISPRi, and proliferation-based CRISPRi screens, we identified a number of both T cell-relevant genes that directly control T cell activation and novel target genes that operate outside of this network cluster, including PPP5C, a protein phosphatase that regulates ERK signaling in cancer cells^44^, but previously unknown to affect T cells. We find that reducing PPP5C expression leads to altered expression of key genes that play a role in T cell biology and function. Through testing variant CRE effects across other cellular functions, we can begin to better understand how common variants across cellular networks affect gene expression and function, and how variants may work together to lead to disease.

Thus, our genomic screens in primary human T cells connect likely causal variants to their putative effects on T cell expression networks and function. These data can be used to propose mechanisms of risk and protection from autoimmune diseases mediated by primary T cells and begin to determine convergent properties of variants, which could be useful for stratifying polygenic risk scores and modifying treatments for individuals to target specific pathways.

## Materials and Methods

### Human Donors

Study protocols were approved by the Benaroya Research Institute IRB under protocol number IRB07109-633. The protocol was conducted according to the principles expressed in the Declaration of Helsinki.

### Cell culture

For MPRA and bulk CRISPRi experiments, eleven donor PBMC were isolated from fresh apheresis leukoreduction packs (BloodWorks) using Ficoll-Paque plus (GE #17-1440-03) (no sequencing of identifiable information), and for single cell CRISPRi screens, we used frozen PBMCs from healthy donors within the BRI Biorepository. CD4 T cells were magnetically isolated (BioLegend #480130) and activated in T cell media (TCM), constituting either 1) X-VIVO 15 (Lonza #04-418Q) supplemented with 25 mM HEPES, 1 mM Sodium Pyruvate, 0.5% Non-essential amino acids, 1% Penicillin Streptomycin, 0.5% L-Glutamine, 5% FBS, 55 mM 2-mercaptoethanol or 2) CTS OpTmizer T cell Expansion medium (ThermoFisher #A1048501) supplemented with 5% FBS, 1% glutamine, 100U/mL PenStrep, 55 mM 2-mercaptoethanol. Cells were activated with human T cell activation beads (Miltenyi #130-091-441) and recombinant human IL-2 (final concentration 100 U/mL; NCI Biological Resources Branch). Lenti-X 293T cells were maintained in DMEM, supplemented with 10% FBS, 1% glutamine, 100U/mL PenStrep, 1 mM sodium pyruvate, 1x MEM non-essential amino acids, and 10 mM HEPES. Cells were passaged every two-three days using trypsin-EDTA for dissociation and kept at a confluency of less than 60%.

### MPRA library transfections

5e7 CD4 T cells were activated for 48-96 hours. Cells were transfected using Neon transfection system (ThermoFisher # MPK10096). In brief, 1e8 activated cells were counted, washed twice in PBS, and resuspended in 1 mL of T buffer and then mixed with 200 μg of MPRA library plasmid from Mouri, et al.^18^ The mixture was divided into 10 reactions and electroporated with the program (2100 volt, 20 ms, 1 pulse). After electroporation, cells were transferred to 50 mL of prewarmed media in T75 flask immediately and then incubated at 37°C for 24 hours. Right before harvesting the cells, transfection efficiency was checked according to flow cytometry, finding 11.9-23.7% GFP+ cells for all seven donors. Cells were then collected by centrifugation, washed once with PBS, collected and frozen at −80°C.

Total RNA was extracted from cells using QIAGEN Maxi RNeasy (QIAGEN, 75162) following the manufacturer’s protocol including the on-column DNase digestion. A second DNase treatment was performed on the purified RNA using 5 μL of Turbo DNase (Life Technologies, AM2238) with buffer, in 750 μL of total volume for 1 hour at 37 °C. The digestion was stopped with the addition of 7.5 μL 10% SDS and 75 μL of 0.5 M EDTA followed by a 5-min incubation at 70 °C. The total reaction was then used for pulldown of GFP mRNA. Water was added to the DNase-digested RNA to bring the total volume to 898 μL to which 900 μL of 20X SSC (Life Technologies, 15557-044), 1800 μL of formamide (Life Technologies, AM9342) and 2 μL of 100 μM biotin-labeled GFP probe (GFP_BiotinCapture_1-3, IDT, Supplementary Table 30) were added and incubated for 2.5 hours at 65 °C. Biotin probes were captured using 400 μL of pre-washed streptavidin beads (Life Technologies, 65001) eluted in 500 μL of 20X SSC. The hybridized RNA/probe bead mixture was agitated on a nutator at room temperature for 15 min. Beads were captured by magnet and washed once with 1× SSC and twice with 0.1× SSC. Elution of RNA was performed by the addition of 25 μL water and heating of the water/bead mixture for 2 min at 70 °C followed by immediate collection of eluent on a magnet. A second elution was performed by incubating the beads with an additional 25 μL of water at 80 °C. A final DNase treatment was performed in 50 μL total volume using 1 μL of Turbo DNase with addition of the buffer incubated for 60 min at 37 °C followed by inactivation with 1 μL of 10% SDS and purification using RNA clean Ampure XP beads (Beckman Coulter, A63987).

First-strand cDNA was synthesized from half of the DNase-treated GFP mRNA with SuperScript III (ThermoFisher, 18080-400) and a primer specific to the 3′ UTR (MPRA_v3_Amp2Sc_R, Supplementary Table 30) using the manufacturer’s recommended protocol, modifying the total reaction volume to 40 μL and performing the elongation step at 47 °C for 80 min. Single-stranded cDNA was purified by Ampure XP and eluted in 30 μL EB.

### MPRA sequencing libraries

To minimize amplification bias during the creation of cDNA tag sequencing libraries, samples were amplified by qPCR to estimate relative concentrations of GFP cDNA using 1 μL of sample in a 10 μL PCR reaction containing 5 μL Q5 NEBNext master mix, 1.7 μL SYBR Green I diluted 1:10,000 (Life Technologies, S-7567) and 0.5 μM of TruSeq_Universal_Adapter and MPRA_Illumina_GFP_F primers (Supplementary Table 30). Samples were amplified with the following qPCR conditions: 95 °C for 20 s, 40 cycles (95 °C for 20 s, 65 °C for 20 s, 72 °C for 30 s), 72 °C for 2 min. The number of cycles for sample amplification was 1−*n* (the number of cycles it took for each sample to pass the threshold) from the qPCR. To add Illumina sequencing adapters, 10 μL of cDNA samples and mpra:minP:gfp plasmid control (diluted to the qPCR cycle range of the samples) were amplified using the reaction conditions from the qPCR scaled to 50 μL, excluding SYBR Green I. Amplified cDNA was SPRI purified and eluted in 40 μL of EB. Individual sequencing barcodes were added to each sample by amplifying the entire 40 μL elution in a 100 μL Q5 NEBNext reaction with 0.5 μM of TruSeq_Universal_Adapter primer and a reverse primer containing a unique 8 bp index (Illumina_Multiplex, Supplementary Table 30) for sample demultiplexing post-sequencing. Samples were amplified at 95 °C for 20 s, six cycles (95 °C for 20 s, 64 °C for 30 s, 72 °C for 30 s), 72 °C for 2 min. Indexed libraries were SPRI purified and pooled according to molar estimates from Agilent TapeStation quantifications. Samples were sequenced using 1 × 30 bp chemistry on a NextSeq 2000 (Illumina).

### Lentivirus production

Protocol was modified from Schmidt et al.^47^. For making lentivirus, Lenti-X 293T cells were seeded in Opti-MEM™ I Reduced Serum Medium, GlutaMAX™ Supplement (ThermoFisher # 51985034) supplemented with 5% FBS, 1 mM sodium pyruvate and 1x MEM-Non-essential amino acids (as cOPTI-MEM) at 4e6 cells per 10 cm petri dish one day before the transfection. Cells were transfected at 80% confluency using 41.4 μL of Lipofectamine 3000 transfection reagent (ThermoFisher #L3000015) in 1250 μL of room temperature plain OPTI-MEM (ThermoFisher # 31985070). 11 ug of transfer plasmid (dCas9-ZIM3-mCherry, Addgene #154473 or sgRNA libraries cloned into CROP-seq-opti, Addgene #106280), 7.5 μg of psPAX2 (Addgene #12260), 3.3 μg of pCMV-VSVG (Addgene #8454) and 36.5 μL of p3000 reagent were added to 1250 μL of room temperature plain OPTI-MEM in a separate tube and mixed by gentle pipetting. The plasmid and Lipofectamine 3000 mixes were combined, mixed by gentle pipetting as 2.5 mL of transfection mixtures, and incubated for 15 minutes at room temperature. Following incubation, 5 mL of medium were removed from the 10 cm dish and 2.5 mL of transfection mixture was added. After 6 hours, the transfection medium was replaced with 15 mL of cOPTI-MEM containing 1x ViralBoost (Alstem Bio #VB100). Lentivirus supernatant was harvested and kept at 4°C for 24 hours after transfection (first harvest) and replaced with 15 mL fresh cOPTI-MEM. The second harvest was done 48 hours after transfection. The two harvests were pooled and spun down at 500 g for 5 minutes at 4°C to clear cell debris. Lenti-X concentrator (Takara Bio #631232) was used to concentrate the virus following the manufacturer’s instruction and resuspend in plain OPTI-MEM at 100-fold concentrated volume of the original volume. Concentrated virus was subsequently aliquoted and frozen at –80°C.

### Generation of CRISPRi libraries

gRNAs libraries for CRISPRi screens were generated through identifying gRNAs that bind in 100 bp (v1 scCRISPRi and proliferation) or 150 bp (v2) windows centered on the variant. gRNAs were obtained within windows using Guidescan ^67^. For scCRISPRi libraries, 10 gRNAs were selected based on the highest sensitivity and specificity scores from GuideScan. gRNA pools were synthesized using Agilent or Twist Biosciences with Gibson Assembly adapters for cloning into the CROP-seq-opti vector (Addgene #106280). These ssOligos were reconstituted to a concentration of 10 nM by dilution with H2O (molecular weights were assessed using NEBioCalculator). For optimal amplification of the library, the following qPCR protocol was implemented. Each reaction (50 µL total) comprised: NEBNext® Q5® Hot Start HiFi PCR Master Mix: 25 µL, 2.5 µL each of sgRNA_libcloning_PCR2_F and sgRNA_libcloning_PCR2_R (5 µM each; Supplementary Table 30), 2.5μL of library at 10 nM, 8.5 µL of SYBR Green (Invitrogen S7563) diluted 10,000 times, and 9 µL of H2O. The reactions were scaled to 12 when making the proliferation library. PCR cycling parameters were determined using NEB Tm calculator: Initial denaturation at 98°C for 30 seconds, 10 cycles (Denaturation at 98°C for 10 seconds, Annealing at 68°C for 30 seconds, Elongation at 65°C for 45 seconds), Final elongation at 65°C for 5 min.

To ensure efficient amplification and specific target enrichment of the library, qPCR monitoring was utilized. The optimal number of cycles was determined when the fluorescence intensity reached between one-third to half of the plateau fluorescence during the exponential phase of amplification. Samples were then removed from the qPCR machine immediately. Following PCR amplification, all reactions were pooled and 1.6x SPRI purification process was used to isolate the amplicons. The purity of the eluted DNA was assessed using Tapestation, while the concentration was quantified using a Qubit fluorometer. Subsequently, the purified amplicons were diluted to a final concentration of 5 ng/uL for subsequent steps.

To linearize the CROP-seq-opti vector, 20 µg of vector was combined with 4 µL of NEB BsmBI v2 restriction enzyme and 8 µL of NEB Buffer r3.1. H2O was added to achieve a total volume of 80 µL. The mixture was incubated overnight at 55°C for restriction digest. The following day, 2 µL of FastAP (ThermoFisher # EF0651) was added to the digestion reaction, and the mixture was incubated at 37°C for 30 min to dephosphorylate the vector ends. Post-incubation, the digested vector underwent a 1.5x SPRI purification to remove enzymes, with an elution step performed in 20 µL of H2O. The concentration of the purified vector was determined using a Qubit fluorometer, and it was subsequently diluted to a final concentration of 250 ng/µL in preparation for downstream cloning procedures.

The vector and insert were combined in a 100 µL Gibson Assembly reaction consisting of 50 µL of NEBuilder® HiFi DNA Assembly Master Mix, 2250 ng (0.4 pmol) linear CROPseq-opti vector, 205 ng library DNA insert: 205 ng (3.36 pmol) and H2O to bring the total volume to 100 µL. The reaction was incubated at 50°C for 1 hour, followed by purification using a 1.5x SPRI method to elute 21 µL of the assembled DNA. The concentration of the assembled DNA was determined using a Qubit fluorometer.

Subsequently, approximately 200 ng of the assembled DNA was transformed into 25 mL NEB 10 beta competent cells via electroporation, following manufacturer’s instructions (2.0 kV, 200 Omega, and 25 μF), and immediately added 975 µl of 37°C NEB 10-beta/Stable Outgrowth Medium. A vector-only control, containing an equal amount of vector DNA, was used as a negative control in the transformation process. Cells were allowed to recover while shaking at 225 RPM for 1 hour at 37°C, then 10 µL of transformed cells was diluted with 90 µL of LB, followed by six 10-fold serial dilutions. Subsequently, 10 µL from each of the six dilutions was streaked onto pre-warmed LB agar plates supplemented with carbenicillin (LB carb+ plates), resulting in dilutions of 1e-3, 1e-4, 1e-5, 1e-6, 1e-7, and 1e-8 of the total electroporated competent cells. The remaining 990 µL of bacterial culture was added to 500 mL of LB broth supplemented with carbenicillin and shaken at 225 RPM at 30°C until reaching an optical density at 600 nm (OD600) of 2.0. Bacteria were then harvested, and plasmids were extracted with ZymoPURE II Plasmid Maxiprep Kit (ZYMO #D4203). The following day, the number of colonies from each dilution of assembled DNA was compared to those from vector-only plates. Successful cloning is indicated only if the efficiency of assembled DNA shows at least a 20-fold increase compared to vector-only controls.

### Single cell CRISPRi screens

5 million CD4+ T cells were thawed and cultured in 5 mL of TCM supplemented with human T cell activation beads for 48 hours. T cells were then infected with a 2-4% (v/v) solution of 100x concentrated dCas9-ZIM3-mCherry lentivirus to introduce the CRISPR interference machinery. 24 hours later, the cells were washed twice with PBS and subsequently infected with an 8% (v/v) solution of 100x concentrated CRISPR-QTL virus, which targets specific variants of interest. 5% of gRNAs contained within the library targeted CD45 as a positive control to evaluate gene suppression efficacy. By day 2 post-library infection, puromycin was added to the culture at a final concentration of 2.0 µg/mL in fresh TCM to select for library transduced cells, and the cell density was adjusted to 0.5 million cells/mL. At 4 days post-infection (4dpi), the cells were resuspended in 20 mL of TCM containing 1.0 µg/mL of puromycin to maintain selection pressure. On days 10 to 12 post-infection, the cells were harvested and divided into six groups.

They were stained with anti-CD3, anti-CD4, and anti-CD45 antibodies, as well as a live/dead dye. Six hashtag antibodies were also used to separate donors and to enable superloading of the 10X controller. Prior to sorting, six groups of cells were pooled, and then sorted. 150K (v1 – 1 donor) and 330K (v2 – 2 donors) mCherry^hi^/GFP^hi^ cells (CD3+/CD4+/live/CD45+ pre-gated) were sorted using a FACS cell sorter. The sorted cells were promptly loaded onto two (v1) or six (v2) channels on the 10x Chromium X controller (10x Genomics) according to the manufacturer’s protocol, with a target capture of 20,000 (library 1) and 12,500 (library 2) cells per channel. Sequencing libraries were generated using the Chromium Next GEM Single Cell 5′ Kit v2 (10x Genomics, #1000265). Gene expression, CRISPR, and feature barcoding libraries were pooled at a 4:1:1 ratio and treated with Illumina Free Adapter Blocking Reagent (Illumina, #20024144). Sequencing of pooled libraries was carried out on a NextSeq 2000 sequencer (Illumina), using a NextSeq P3 flowcell (Illumina) for v1 or sequenced on a Nova-seq X Plus 25B flow cell. Basecalls were processed to FASTQs on BaseSpace (Illumina).

### Bulk CRISPRi screen for T cell proliferation

6e7 CD4+ T cells were activated in 30 mL of TCM for 24 hours. T cells were then infected with 2% v/v 100x concentrated dCas9-ZIM3-mCherry lentivirus. 24 hours later, the cells were infected again with 0.25% v/v 100x concentrated GW2 library lentivirus (MOI 0.5∼1). At 1 day post-infection (dpi), cells were counted to ensure there were at least 30e6 live cells. 15e6 cells were collected as time zero control (day 2) of proliferation screens. For the remaining cells, fresh Th0 media (TCM with recombinant human IL-2 (final concentration 500 U/mL)) and puromycin (final concentration 2.5 μg/mL) was added to bring cells to 1e6 cells/mL. At 2 dpi, cells were harvested, spun down and resuspended at 0.5e6 cells/mL in Th0 media. Cells were maintained between 0.5-1e6 cells/mL until 10 dpi, when cells were harvested and live ZIM3-mCherry+ cells were FACS sorted (typically ∼10-30% of total cells). Sorted ZIM3-mCherry+ cells were maintained at 0.5e6-1e6 cells/mL in Th0 media until day 21. At least 15M cells were then collected per donor and stored at –80°C until genomic DNA extraction.

### Bulk CRISPRi screen sequencing library preparation

Genomic DNA from cells (in 10M cell increments) were resuspended in 50 μl ChIP lysis buffer (1% SDS, 10 mM EDTA, in 50 mM Tris-HCl, pH 8.1) and pipetted up and down. Lysed cells were transfered to a 96-well plate or 8-well strip, then incubated at 65°C for 10 min. Then sample was cooled to 37°C, and 1 μl Rnase cocktail (Ambion, AM2286) was added, again mixed by pipetting and spun down followed by incubation at 37°C for 30 min. 5 μl proteinase K (NEB, P8107) was added and the sample was mixed by pipetting. The sample was then incubated at 37°C for 2 hours, then at 95°C for 20 min to denature the proteinase K. To isolate genomic DNA, we added 36.4μl Ampure XP to the sample, mixed thoroughly by pipetting, incubated for 5 min, and used a magnet to isolate genomic beads from the lysed sample. We pipetted off the supernatant and washed the sample 3 times with 80% ethanol while on the magnet. After drying the pellet for 5 min, gDNA was then eluted in 45 μl ddH2O, yielding on average 3-6 pg per cell.

We then used 0.6 μg of genomic DNA from each sample for qPCR to determine optimal PCR cycle number for library preparation. Each 10 μL qPCR reaction contained 5 μL of NEBNext Q5 Hotstart HiFi PCR master mix (NEB # M0543L), FwdInnerSeq and RevInnerSeq at 500 nM each (final concentration; Supplementary Table 30), 0.6 μg of genomic DNA, 1.7 μL of SYBR (diluted 1/10000), and added water to have a final volume of 10μL. Once the optimal PCR cycle was determined, 50 μL PCR reactions were used to amplify the amplicon, and the number of reactions were scaled to the total available gDNA. The PCR reactions from each sample were pooled, and 50 μL was taken for amplicon purification. Amplicons were purified using a two-step Ampure XP method. 0.65x Ampure XP was added to the sample, followed by mixing via pipet, and incubating for 5 min. The sample was applied to the magnet and the supernatant was isolated for further purification. To the supernatant, an additional 1.0x Ampure XP was added to the sample to capture PCR amplicon. The sample was incubated for 5 minutes, and then magnetic beads were captured by magnet. The supernatant was removed and discarded, and the captured beads were washed 3 times with 80% ethanol. The sample was dried for 5 min and eluted with 25 μL of H2O. Samples were analyzed on a TapeStation to assess amplicon purity and size estimation, and the concentration of amplicon DNA was measured using Qubit. Samples were pooled based on the concentration of the specific target amplicon percentage and sequenced on a Nextseq 2000 with custom read1 primer hU6_R1 and custom index primer sgPuro_I.

### Bulk RNAseq

1e7 frozen CD4+ T cells were thawed and activated using human T cell activation beads in 10 mL of TCM. 24 hours later, the T cells were infected with a 10-15% v/v solution of 100x concentrated dCas9-ZIM3-mCherry lentivirus, facilitating the introduction of the CRISPR interference machinery. 24 hours after the CRISPRi infection, the cells were further infected with a 10-15% v/v solution of 100x concentrated gRNA virus designed to target specific variants of interest (Supplementary Table 30 for gRNA sequences). As a positive control we utilized a CD45 sgRNA to assess the efficacy of gene suppression. On day 2 post-guide infection, puromycin was added into the culture at a final concentration of 2.0 µg/mL in fresh TCM to select for transduced cells and adjust the cell density to 0.5 million cells/mL. On day 4, the puromycin concentration was reduced to 1.0 µg/mL to maintain selection. Cells were expanded for an additional 3-4 days. One day prior to cell sorting, flow cytometry analysis was performed to confirm the efficiency of CD45 knockdown (>85%) in mCherryhi / GFPhi cell populations. On day 7 or 8, a flow sorter was utilized to isolate at least 300k cells exhibiting high expression levels of mCherryhi / GFPhi. Post-sorting, cells were centrifuged to remove supernatant and immediately lysed using trizol reagent. Cell lysates were subsequently frozen at –80°C prior to RNA extraction. RNA extraction was carried out using Direct-zol RNA Microprep (ZYMO # R2063), following the manufacturer’s instructions, to obtain RNA for bulk RNA-seq. Total RNA was added to reaction buffer from the SMART-Seq v4 Ultra Low Input RNA Kit for Sequencing (Takara, #634891), and reverse transcription was performed followed by PCR amplification to generate full length amplified cDNA. Sequencing libraries were constructed using the NexteraXT DNA library preparation kit with unique dual indexes (Illumina, FC-131-1096) to generate Illumina-compatible barcoded libraries. Libraries were pooled and quantified using a Qubit® Fluorometer (Life Technologies). Sequencing of pooled libraries was carried out on a NextSeq 2000 sequencer (Illumina) with paired-end 59-base reads, using a NextSeq P2 sequencing kit (Illumina) with a target depth of 5 million reads per sample.

## Analysis Methods

### MPRA data processing

MPRA data for primary T cells and Jurkat cells^18^ were analyzed using MPRASuite (https://github.com/tewhey-lab/MPRASuite), which comprises of MPRAmatch (barcode/oligo pairing), MPRAcount (tag assignment and counting), and MPRAmodel (activity and allelic effect estimates). MPRAmodel employs DESeq2^68^ for normalization and statistical analysis. The paired-end reads were merged into single amplicons using Flash^69^ to identify oligo/barcode pairings. Genomic DNA sequences and barcodes were extracted and mapped back to the oligo design file using minimap2^70^. Alignments with more than a 5% error rate were discarded, retaining only high-quality alignments, which were compiled into a lookup table with their matching barcodes. After sequencing the RNA and plasmid libraries, 19 bp barcode tags were assigned to oligos from the lookup table, and oligo counts were aggregated by summing across all barcodes. Variants with fewer than 30 DNA counts or with zero RNA counts in oligos for either allele were excluded from further analysis.

We normalized the MPRA samples using MPRAmodel, where the library counts for DNA plasmid replicates with RNA Jurkat cells (3 replicates) and separately with primary T cells (7 replicates) were normalized across samples using the “summit-shift” method, which shifts the log2(fold change) density peak to line up with 0 and centers the RNA/DNA ratio distribution of each sample around 1, effectively assuming that most genomic elements in the MPRA are minimally active (which we anticipate is true for this library because most variants included for each association are not causal).

To define active elements, significant differences in counts between plasmid (DNA) and Jurkat and primary T cell-types (RNA) were determined using a negative binomial generalized linear model (GLM) within DESeq2. This model incorporated independent dispersion estimates for each cell-type and library. Activity of genomic elements was quantified in log2(fold-change) units comparing counts from RNA versus DNA, using a contrast in DESeq2’s design matrix to compare treatment conditions and Wald’s test to assess significance. Multiple hypothesis testing within each cell-type and library was corrected using Bonferroni’s method. Post-MPRAmodel analysis, a filter for a plasmid count of >20 was employed for the primary T cell MPRA data for all downstream analyses. This brought the total number of variants from 18326 to 18182.

### Defining MPRA activity thresholds using a DHS enrichment grid search

To empirically identify an activity cutoff for primary T cell putative CREs and emVars, we performed a grid search to determine the enrichment of active MPRA elements within primary T cell DHS sites for both primary T cell and unstimulated Jurkat MPRAs similar to the epigenetic DHS enrichment analysis above. Briefly, we used a grid search for active element enrichment within CD4 T cell DHS sites and obtained a *P* value based on different activity thresholds log2(fold-change [FC]) and activity *P* values (log10(*P* Adj. BH)). The grid search is filled with variant level enrichment (*E*_*v*_) values and a *P* value (using a two-by-two contingency table with two-sided Fishers exact test) for the set of pCREs created using the values on each axis (Supplementary Tables 32 and 33). The equation for enrichment is shown, represented as a risk ratio:

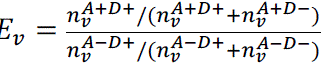

Here, *n_v_* is the number of variants. Superscript *A* + indicates that it is an active element while *A* − indicates that the element is inactive, and superscript *D* + again refers to whether the variant is within a DHS site (*D* + if emVar, *D* − otherwise).

For each activity logFC and logFC *P* value cutoff, color indicates the fold enrichment for T cell DHS sites and the dot size is the sample size of the true positive (active elements in a DHS peak). Each axis indicates the tested cutoffs for log2(FC) and log10(*P* Adj. BH). As seen in the bottom row of the grid search plots, the primary T cell active elements have a larger starting sample size (maximum 449 elements). We created a log2FC cutoff for primary T cells to increase the active element DHS enrichment while maintaining a sufficient sample size and enrichment *P* value. Because of the initial low sample size of active elements within the Jurkat MPRAs and the high initial enrichment *P* values for T cell DHS sites, we chose to include all active elements regardless of activity for this cell type.

### Calling emVars

Allelic effects were estimated in Δ log2(fold-change) units, comparing RNA counts between the Alternate (Alt) and Reference (Ref) oligos. This was achieved by including a contrast in DESeq2’s design matrix for alleles (Alt vs Ref) alongside treatment conditions followed by Wald’s test for significance and FDR estimation using Benjamini and Hochberg’s method. We then divided the variants into three categories, emVars (expression modulating variants), putative CREs, and no activity variants. For primary T cell MPRAs, a putative CRE must have an expression Bonferroni (BF) adjusted log_10_ *P* value greater than to 2 and an expression log_2_(FC) of greater than or equal to 1 for either allele A or B. emVars must have an expression BF adjusted log_10_(*P* value) greater than to 2, a skew log_10_ (FDR *P* value) greater than or equal to 1, and an expression log fold change of greater than or equal to 1. No activity variants are all the other variants.

### Epigenetic enrichment of emVars and pCREs

DHS data across 733 samples was obtained from Meuleman *et al.*^71^. Pre-processed DHS peaks lifted to hg38 were downloaded from https://zenodo.org/record/3838751#.X_IA7-lKg6U. For each of 261 unique cell types, DHS peaks were merged across replicates of the same cell type. To assess enrichment between DHS sites for each cell type and MPRA data, we counted the number of MPRA variants with regulatory activity (i.e., putative CRE) overlapping the genomic interval bed file for each cell type. We then constructed a 2 x 2 contingency table based on whether the MPRA SNP showed regulatory activity (i.e., putative CRE) or had no regulatory activity, and whether the SNP intersected a genomic interval or not. *P* values were calculated based on a two-sided Fisher’s exact test (Supplementary Table 34). Multiple testing correction was performed using Bonferroni correction by taking an alpha of 0.05 divided by the number of unique cell types tested (n = 261).

Histone ChIP-seq data were obtained from the ENCODE project for all available human CD4-positive alpha-beta T cell samples. For each cell type, all available ChIP-seq peak call sets for H3K4me1, H3K4me3, H3K27ac, and H3K36me3 aligned to hg19 were downloaded in bed file format. If multiple replicates were available, peaks call sets were merged using the merge function in BEDTools v2.26.0^72^. Human CAGE-based enhancer sequences were downloaded from https://fantom.gsc.riken.jp/5/datafiles/latest/extra/Enhancers/ in bed file format. chromHMM annotations in the 18-chromatin state model were obtained for primary T cells from peripheral blood (E034) from the Roadmap Epigenomics Project (https://egg2.wustl.edu/roadmap/data/byFileType/chromhmmSegmentations/ChmmModels/core_K27ac/jointModel/final/) in bed file format. To assess enrichment between each of these genomic datasets and MPRA data, we counted the number of emVar SNPs overlapping the genomic interval bed file. We then constructed a 2 x 2 contingency table based on whether the SNP showed MPRA activity (i.e., emVar) or had no MPRA activity, and whether the SNP intersected a genomic interval or not. *P* values were calculated based on a two-sided Fisher’s exact test. Multiple testing correction was performed using Bonferroni correction by taking an alpha of 0.05 divided by the number of genomic annotations tested.

### ATAC-seq skew and QTL enrichment of emVars

To assess enrichment of MPRA pCREs and emVars at sites of allelic skew in open chromatin regions, we downloaded significant allelic skew SNPs from Calderon *et al.*^13^. For this calculation, we used all stimulated hematopoietic cell types. For SNPs with evidence of skew across multiple samples or cell types, we summed the reference and alternate allele counts for that SNP across the cell types. We used a two-sided 2-sample test for equality of proportions with continuity correction to compare the proportion of variants with ATAC-seq allelic skew which are pCREs (χ^2^=61.9, df=1) or emVars (X^2^=67.249, df=1) to those which are no activity variants (Fig. S4D). We also correlated ATAC-seq allelic bias to the MPRA allelic bias (β=0.9302, SE=0.7061, F=5.428, df=23, Fig. S4E). To assess enrichment of MPRA SNPs at ATAC-QTLs, we downloaded primary T cell ATAC-QTLs from Gate, *et al.*^73^. Similarly, we used a two-sided 2-sample test for equality of proportions with continuity correction to compare the proportion of ATAC-QTL variants which are pCREs (χ^2^=20.18, df=1) or emVars (X^2^=5.5258, df=1) to those which are no activity variants (Fig. S4F). We also correlated caQTL effect size (beta) to the MPRA allelic bias (β=0.8192, SE=0.4906, F=0.2974, df=12, Fig. S4G).

### *In silico* predictions of effect of regulatory activity of emVars

We applied deltaSVM v1.3 to predict the effect of MPRA SNPs on regulatory activity^74^. This method uses a classifier (gkm-SVM) to encode cell-specific regulatory sequence vocabularies, and then subsequently deltaSVM quantifies the effect of a SNP as the change in gkm-SVM score). We used the previously calculated deltaSVM score for the variants of this MPRA library calculated for Mouri et al.^18^ using downloaded pre-computed gkm-SVM weights derived from ENCODE2 enhancers in naïve CD4 T cells (http://www.beerlab.org/deltasvm_models/downloads/) and deltasvm.pl from the software developers. Default parameters were used in calculating deltaSVM scores. We correlated the deltaSVM scores with MPRA allelic bias (β=0.07440, SE=1.557, F=2.638, df=484, Fig. S4H).

### Transcription factor enrichment of emVars and pCREs

To replicate the transcription factor enrichment analysis in Mouri et al.^18^, we applied motifbreakR v2.14.2 to predict effect of each MPRA variant on TF binding^75^. For each variant in the MPRA data, we calculated TF binding scores for the reference and alternate alleles. We used the sum of log probabilities approach in motifbreakR, applied to all TF position-weighted matrices in HOCOMOCO v10^76^. All MPRA SNPs with a difference in TF binding scores between the reference and alternate alleles at *P* < 1×10^−5^ were considered to be significant. Additionally, only variants which overlapped with chip-seq data obtained from ENCODE were used (https://hgdownload.cse.ucsc.edu/goldenpath/hg19/encodeDCC/wgEncodeRegTfbsClustered/wgEncodeRegTfbsClusteredV3.bed.gz). We used a two-sided 2-sample test for equality of proportions with continuity correction to compare the proportion of variants with motifbreakR scores which are pCREs (χ^2^=86.712, df=1) or emVars (X^2^=27.436, df=1) to those which are no activity variants (Fig. S4I). We also correlated motifbreakR scores to the MPRA allelic bias of emVars (β=0.20343, SE=0.3882, F=31.89, df=7, Fig. S4J).

### emVar enrichment calculations for fine-mapped SNPs

To determine enrichment of emVars for fine-mapped variants, the HLA region and specific loci with abnormal LD pattern, for which fine mapping performs poorly, were eliminated. The lead SNPs for the loci with abnormal LD patterns are 22:50966914:T:C, 3:105558837:G:A,12:9905851:A:C, 13:40745693:G:A, 16:1073552:A:G, 17:38775150:C:T, 17:44073889:A:G, 18:12830538:G:A, 2:100764087:T:G,21:36488822:T:C, 21:45621817:A:G, 6:127457260:A:G, 6:130348257:C:T, 7:116895163:G:A, 7:51028987:T:A, 2:204592021:G:A, and 14:75961511:C:T. The coordinates for the HLA region used are chr6:29691116-33054976. This filter reduced the total number of variants in the analysis from 18182 to 14514.

To calculate enrichment of MPRA emVars in PICS fine-mapped SNPs, we determined for each MPRA variant whether it was a PICS statistically fine-mapped SNP (i.e., had a PICS probability greater than a given threshold) and whether the fine-mapped SNP showed MPRA activity (i.e., emVar). From this, we constructed 2 x 2 contingency tables. We then calculated variant-level enrichment (*E*_*v*_), represented as a risk ratio:

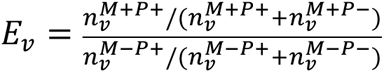

Here, *n_v_* refers to the number of variants. Superscript *M* refers to whether the variant is an emVar (*M* + if emVar, *M* − otherwise). Superscript Prefers to whether the SNP is a statistically fine-mapped SNP by PICS (P + if PICS fine-mapped, P − otherwise). We can then construct a 2 x 2 contingency table with values n^M+P+^, n^M+P-^, n^M-P+^, and n^M-P-^. Statistical significance was calculated using a two-sided Fisher’s exact test based off this 2 x 2 contingency table.

We similarly calculated enrichment of MPRA variants overlapping T cell DHS sites in PICS statistically fine-mapped SNPs. We determined for each MPRA variant whether it directly overlapped a T cell DHS site and whether it was a PICS fine-mapped SNP (i.e., had a PICS probability greater than a given threshold). We then calculated variant-level enrichment (*E*_*v*_) represented as a risk ratio:

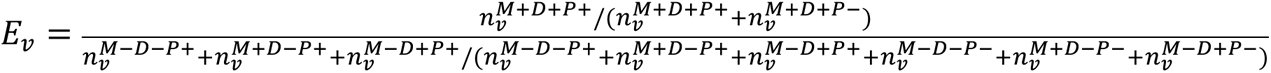

Here, *n_v_* refers to the number of GWAS loci. Superscript *M* refers to whether the variant is an emVar (*M* + if emVar, *M* − otherwise). Superscript P refers to whether the SNP is a statistically fine-mapped SNP by PICS (P + if PICS fine-mapped, P − otherwise). Superscript *D* refers to whether the variant is within a DHS site (*D* + if emVar, *D* − otherwise) and superscript *M* refers to whether the variant is an emVar (*M* + if emVar, *M* − otherwise). We can then construct a 2 x 2 contingency table with values 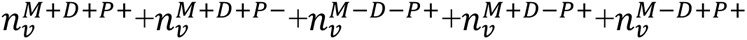. and 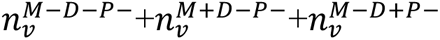. *P* values were calculated using a two-sided Fisher’s exact test. Using the same process, we calculated enrichment of MPRA emVars in UK biobank fine mapping. The UK biobank data did not overlap completely with our variants in the MPRA library.

We also calculated sensitivity and specificity of the MPRA using PICS fine-mapping as the benchmark “ground truth.” To do this, we tabulated for each locus, if there was a fine-mapped SNP (i.e., in X% credible set) and whether the fine-mapped SNP showed MPRA activity (i.e., emVar). We can then calculate sensitivity at the locus level (*SE_l_*):

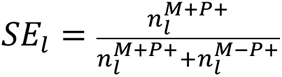

And specificity (*S*P*_l_*):

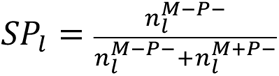

Here, *n_l_* refers to the number of GWAS loci. Superscript *M* refers to whether the variant is an emVar (*M* + if emVar, *M* − otherwise). Superscript P refers to whether the SNP is a statistically fine-mapped SNP by PICS (P + if PICS fine-mapped, P − otherwise).

### MPRA expression in elements that perturb TF motifs

To assess TF binding to MPRA-tested variants, we ran variants in the MPRA library through the motifbreakR v2.14.2^75^ algorithm to identify TFs which have a differential allele-specific binding strength to specific elements. In contrast to the TF analysis above which replicated Mouri et al.^18^, we used a 1e-4 p-value cutoff and Hocomoco v11^76^ to generate these motifbreakR results. Using MPRA data, we compared the differential expression between alleles of variant (based on the LogSkew value) that disrupts a TF motif (filtering for those that have at least five MPRA variants disrupting a TF motif) to those that did not using a two-sided *t* test. We checked histograms and QQ plots of LogSkew of variants that disrupts TFs to confirm that LogSkew is normally distributed.

MotifbreakR gives allele-specific binding scores which include some variants in which the reference allele binds more strongly to a motif and some which bind less (or disrupt the motif). For those reference alleles that bound to the TF more strongly, we flipped the reference and alternate alleles (flipping the sign of LogSkew) so that our test would examine all variant alleles that disrupt TF binding. Since we flipped some of the alleles of variants that bound to a given TF, we chose to flip an equal proportion of those which did not bind to that TF, repeating this test 100 times for randomly selected non-TF binding variants. Any TFs that had a *P* value of >0.05 in any of the 100 tests is considered non-significant and grayed out in the volcano plots (Supplementary Fig. 6). Additionally, we filtered out TFs according to their expression in either Jurkat or primary T cells from RNA-seq data (≥1 TPM)^77,78^.

Next, in primary T-cells we examined variants that we PICS fine-mapped for each disease (MS, RA, T1D, psoriasis, IBD, UC, and Crohn’s). Focusing on variants associated with each disease, we ran the same test as above to determine TFs disrupted by disease-associated variants that also correlate with allele-specific expression again filtering for TFs that have at least 5 disease-associated variants disrupting the TF’s motif (Supplementary Table 35). For the top 5 TFs by *P*-value for positive and negative effect size for each disease, we collated the data for all the diseases, hierarchically clustered the data (Euclidean distance), and created a heatmap of all the TFs and with the *P* values and effect sizes for their respective diseases. We then compared data from both the Jurkat and primary T cell MPRAs to see the differences in activity of variants which disrupted TF binding, plotting their effect sizes and *P* values for TFs at least expressed in one cell type.

### V2G Network analysis

We connected variants to target genes using Open Targets Genetics V2G. To do this, we first utilized the Genopyc tool to convert rsID to variant ID^79^. Subsequently, these variant IDs were associated with genes using the Open Targets V2g pipeline^19^ with the otargen package in R^80^. The V2G output underwent processing using pandas and polars to filter out non-T cell-related terms for QTL and capture Hi-C targets. Following this, the V2G output was filtered for genes expressed in T cells (TPM > 1) using the reference data from the DICE database^77^. This systematic approach was used to generate the background and foreground files for further analysis. Genes from the V2G list were input into STRING and networks were built according to all interaction sources (Textmining, Experiments, Databases, Co-expression, Neighborhood, Gene Fusion, Co-ocurrence). Network in Fig. 3 had a foreground of Open Targets V2G of primary T cell emVars in T cell DHS sites and a background of V2G for all MPRA-tested variants in T cell DHS sites, and clusters were made with MCL clustering with an inflation parameter of 2.2. This cluster had 667 node genes and 3,125 edges (PPI enrichment *P*-value=1e-16). Supplementary Fig. 7 had a foreground of V2G genes of Jurkat emVars in T cell DHS sites and a background of V2G for all MPRA tested variants in T cell DHS sites, and clusters were made with MCL clustering with an inflation parameter of 2.2. This cluster had 336 gene nodes and 970 edges (PPI enrichment *P*-value=2.2e-16). Fig. 4f had a foreground of genes identified in single cell CRISPRi screens that were identified as local targets of 56 emVars in T cell DHS and the background was all potential target genes within 1 Mb of each of the 56 emVars in T cell DHS that were expressed in T cells, and clusters were made with MCL clustering with an inflation parameter of 2.0. This cluster had 55 node genes and 42 edges (PPI enrichment *P*-value=0.398). Figure 5c was generated with a foreground of V2G genes of emVars in T cell DHS that were hits within the proliferation screen (FDR < 0.1), and the background was all V2G genes connected to the 56 emVars in T cell DHS that were tested within the proliferation screen, and clusters were made with MCL clustering with an inflation parameter of 2.0. This cluster had 142 node genes with 257 edges (PPI enrichment *P*-value=4.29e-09).

### Single-cell CRISPRi screen processing

Demultiplexed reads were processed using Cell Ranger software version 7.1.0 (10x Genomics) in combination with the GRCh38-2020-A reference genome (10x Genomics). The cellranger ‘mult’ function was used to quantify gene expression, feature bar code counts and CRISPR guide RNA (gRNA) counts. The resulting gene expression, feature barcode and CRISPR gRNA count matrices were imported into Seurat version 5.0.0^81^. Standard preprocessing steps were applied, including log-normalization of the gene expression data and identification of highly variable genes. The Seurat pipeline was used for scaling the data and regressing out unwanted sources of variation. Principal Component Analysis (PCA) was performed to reduce the dimensionality of the dataset, followed by clustering using the Louvain algorithm with a resolution parameter optimized based on the dataset size and biological context. UMAP (Uniform Manifold Approximation and Projection) was used for visualizing the clusters in a two-dimensional space, facilitating the identification of distinct cell populations. The feature barcode matrix was processed in Seurat for normalization and quality control. A manual cutoff based on the hashtag counts was applied to assign donor identities to individual cells.

### Single-cell CRISPR Screen Data Analysis

This clustering information provided essential context for interpreting the effects of gRNA perturbations in different cell populations. Sceptre version 0.10.0^20,35,82^, an R package designed for analyzing single-cell CRISPR screen data, was employed to evaluate the impact of CRISPR perturbations on gene expression across the identified clusters. gRNA integration was done in low moi mode, and Sceptre’s statistical models were fitted incorporating cluster and donor identities to improve the precision of the effect size estimates.

### Bulk CRISPRi screen analysis

Proliferation screen FASTQs were processed using MAGeCK RRA (v0.5.9.5)^45^ to create gRNA readcount matrices using mageck –count. Differential gRNA enrichment was conducted using mageck test with –-control-sgrna and –-paired flags. Results were analyzed using MAGeCKFlute (2.2.0)^83^.

### Bulk RNA-seq

Base calls were processed to FASTQs on BaseSpace (Illumina). A base call quality-trimming step was applied to remove low-confidence base calls from the ends of reads using trimmomatic (0.38)^84^. The FASTQs were aligned to the Ensemble Human genome assembly version GRCh38.91, using STAR (2.7.11a)^85^ and gene counts were generated using htseq-count (2.0.2)^86^. QC and metrics analysis were performed using fastQC (0.12.1) and the Picard family of tools (3.1.0). Gene count matrices were then input into DESeq2 (1.38.3)^68^. Normalization and mean-variance modeling were conducted through estimateSizeFactors and DESeq. Differential expression of PPP5C TSS gRNA, rs62136101 gRNA, and several other variant targets were contrasted against the NT gRNA. *P*-values were calculated for differential expression within DESeq2 using a Wald test, with Benjamini and Hochberg method for adjusting *P*-values for multiple comparisons. Each contrast was rankordered by *P*-value and written into a separate .csv file.

## Acknowledgments

We thank Drs. Hilary Finucane and Jacob Ulirsch for UKBB fine-mapping data. We thank Dr. Susan Kales for plasmid preparation for MPRAs. We thank Dr. Virginia Green, Dr. Adam Lacy Hulbert, Dr. Jessica Hamerman, Dr. Karen Cerosaletti, Dr. Matthew Dufort, and Matthew Lawrence for critical review of the manuscript. We thank Dr. Naresh Doni Jayavelu and Dr. Matt Altman for advice on network approaches. We acknowledge the Benaroya Research Institute’s Innovation fund, the BRI Clinical Core, the BRI Genomics Core, and the BRI CATA Core, which enabled completion of this work. We thank the M.J. Murdock Charitable Trust for the purchase of scientific instrumentation, which enabled completion of the experiments within this manuscript.

## Author contributions

Conceptualization: C.H.H., J.P.R.

Methodology: C.H.H., M.D., M.M., A.M., H.C., F.C., T.M., J.S., R.T., J.P.R.

Investigation: C.H.H., M.D., M.M., A.M., L.N., S.H.

Visualization: M.D., S.P., H.S., A.M., J.P.R.

Funding acquisition: R.T., J.P.R.

Project administration: C.H.H., J.P.R.

Supervision: H.D., J.H.B., J.S., C.G.B., M.G., R.T., J.P.R.

Writing – original draft: J.P.R.

## Funding

National Institutes of Health grant DP2 AI183504 (J.P.R.)

National Institutes of Health grant U01 AI176320 (J.H.B., J.P.R.)

National Institutes of Health grant K22 AI153648 (J.P.R.)

Crohn’s & Colitis Foundation grant 1158945 (J.P.R.)

National Institutes of Health grant R01AI151051 (R.T.)

National Institutes of Health grant R35HG011329 (R.T.)

National Institutes of Health grant K08 AG086591 (M.H.G.)

Michael Smith Health Research British Columbia Scholar (C.G.D.)

Natural Sciences and Engineering Research Council of Canada, Banting Postdoctoral Fellowship (T.A.M.)

## Data and materials availability

All data, code, and materials used in the analysis is available upon request or publication of the manuscript.

