## Supplementary Information for "Linking candidate causal autoimmune variants to T cell networks using genetic and epigenetic screens in primary human T cells"

**Affiliations:**

**Contents:**

Supplementary Text

Supplementary Figs. 1 - 11

Captions for Supplemental Tables 1 - 35

### Supplementary Text

Primary T cell emVars are enriched for regulatory region features

We observed primary T cell emVars were most highly enriched in transcription start site (TSS) regions and distal enhancers (Supplementary Fig. 3a and b; Supplementary Table 3), with particularly high enrichment at regions marked by H3K4me3, CAGE and DNase hypersensitivity sites (DHS) (Supplementary Fig. 3c; Supplementary Tables 4 and 5), consistent with many emVars altering regulatory element activity. emVars were also more likely to have allelic bias in ATAC-seq data from hematopoietic cell types and to be a chromatin accessibility quantitative trait locus (caQTL) as compared to MPRA variants with no activity, and emVar allelic effects were correlated with allelic bias and QTL directionality from these data (Supplementary Fig. 3d-g). Consistent with emVars disrupting regulatory element activity and chromatin accessibility, we found that their allelic effects were correlated with computationally predicted allelic effects ("delta SVM") in CD4 T cell enhancer elements, with most emVars showing directional concordance with the delta SVM score (Supplementary Fig. 3h). emVars were also much more likely to perturb a transcription factor (TF) motif (according to position weight matrices) when compared to all variants tested in the MPRA assay, with predicted TF binding also correlating strongly with the observed MPRA allelic bias for emVars (Supplementary Fig. 3i-j). Therefore, primary T cell emVars enrich in regulatory regions and for variants that have orthogonal regulatory phenotypes and allele-specific activities.

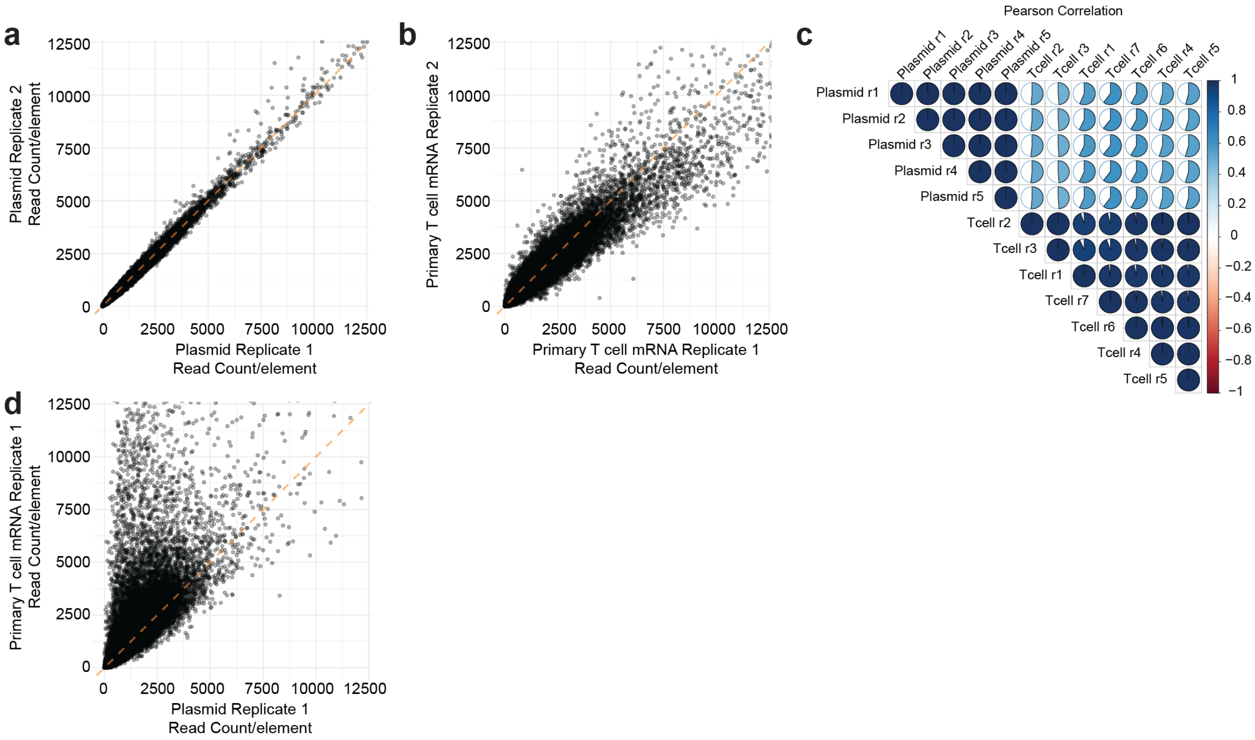


**Supplementary Fig. 1.**

**Primary T cell MPRAs have robust replication. a,** Scatterplot showing correlation between two replicates of plasmid barcode prevalence according to normalized read counts. **b,** Scatterplot showing correlation between two replicates of primary T cell barcode prevalence according to normalized read counts. **c,** Pie charts indicate pairwise Pearson correlation between plasmid and primary T cell replicates. **d,** Scatterplot showing correlation between primary T cell (y-axis) versus plasmid (x-axis) barcode prevalence according to normalized read counts.

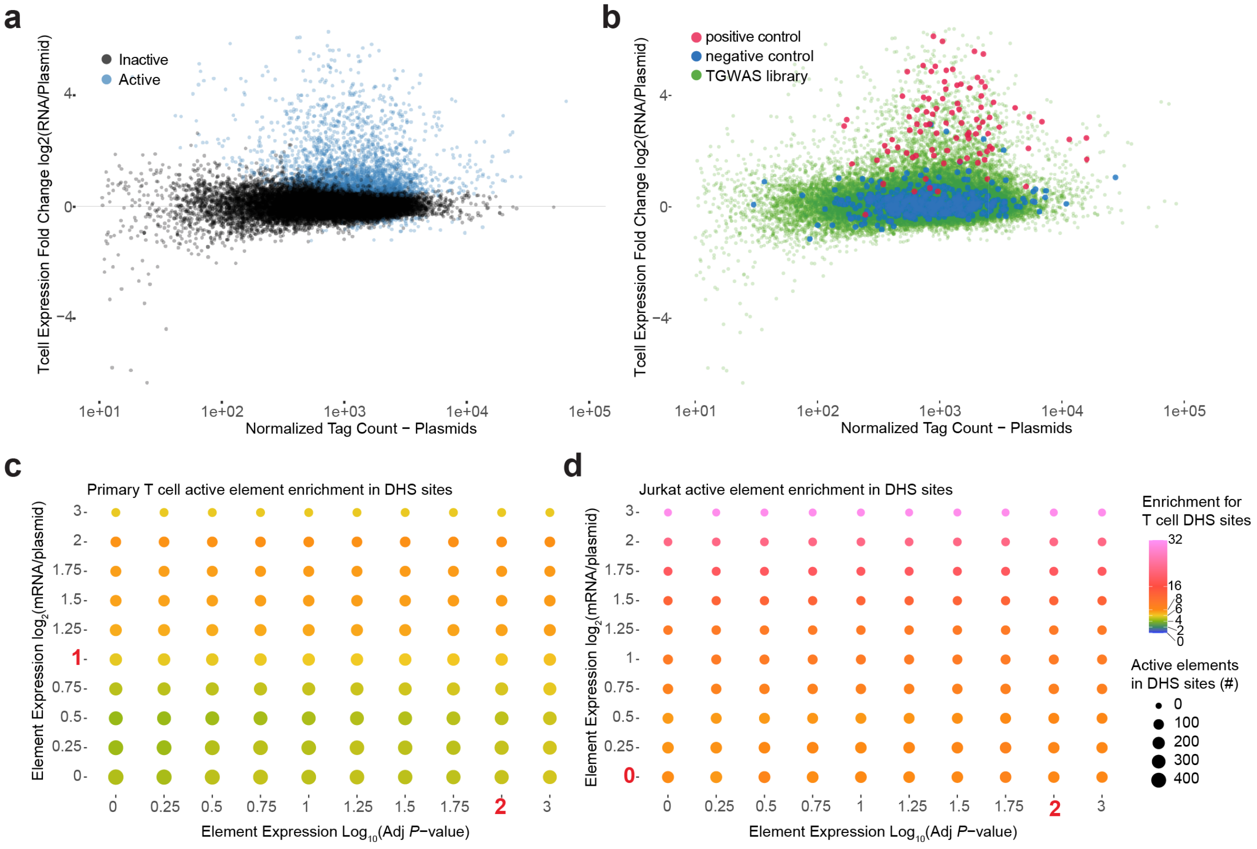


Grid search. Dots represent enrichment (color) and number of elements (size) at each cutoff of expression fold-change (y-axis) and expression FDR value (x-axis).

**Supplementary Fig. 2.**

**Identification of active putative cis-regulatory regions with primary T cell MPRAs. a,** Scatterplot showing normalized tag count (x-axis) by expression fold change of barcode counts in RNA versus plasmid libraries. **b,** Same scatterplot as (a) but indicating spiked in positive (red) and negative (blue) controls. Variant library is indicated in green. **c and d,** Grid search assessing enrichment of expressed elements for primary T cell DHS sites at the given thresholds of expression fold-change ($\log_2$ mRNA/plasmid; y-axis) and expression significance ($\log_{10}$ adjusted *P* value for element expression over baseline; x-axis). In red are the chosen cutoffs for calling putative CREs for primary T cell (c) and Jurkat cell (d) MPRAs. *P* values for (c) and (d) are calculated using a Wald test and adjusted using the Benjamini and Hochberg method.

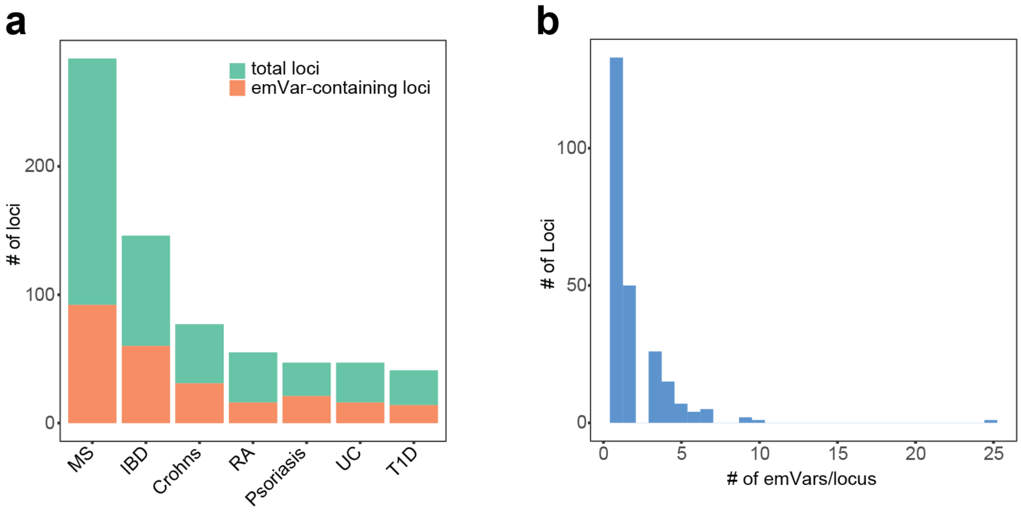


**Supplementary Fig. 3.**

**Primary T cell MPRA prioritizes variants in hundreds of loci. a,** Total number of GWAS loci tested (green) and number of loci with at least one emVar identified (orange) for each disease GWAS. **b,** Histogram of the number of emVars within each GWAS locus.

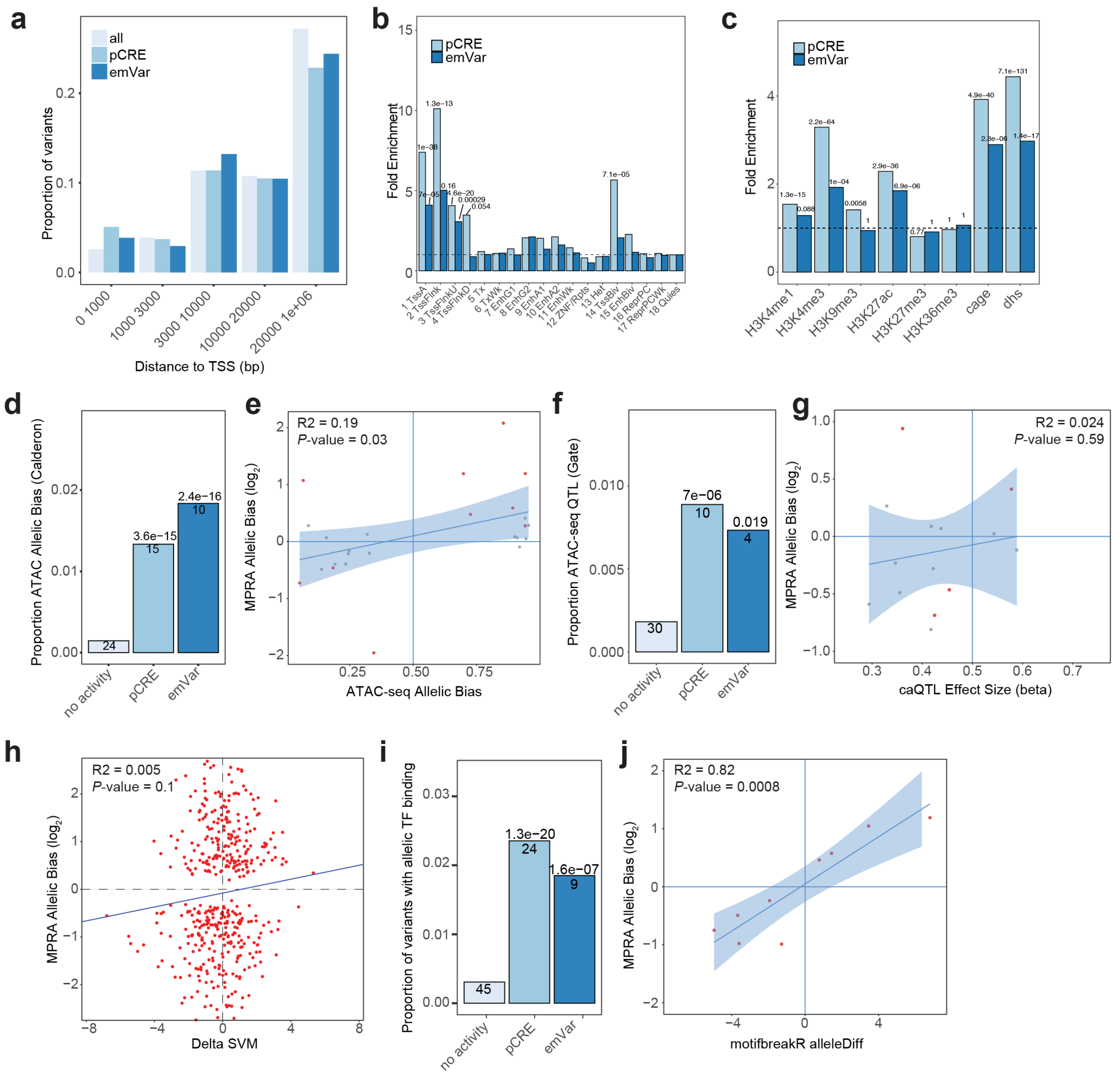


**Supplementary Fig. 4.**

**Variant locations relative to cis-regulatory features. a,** Location relative to TSSs of all MPRA tested variants, active elements (pCRE), and emVars. **b,** Enrichment of variants within pCREs (light blue) and emVars (dark blue) within chromHMM-defined genomic regions in human T cells. (p value Bonferroni-corrected for 36 independent tests). **c,** Functional enrichment of variants within pCREs and emVars (nominal *P* value threshold of 0.05 Bonferroni-corrected for 8 independent tests). **d,** Proportion of inactive element and pCRE variants and emVars that have allelic bias in ATAC-seq. **e,** Scatter plot comparing MPRA log2 allelic bias (y-axis) with allelic bias in ATAC-seq from hematopoietic cells (x-axis)[13]. Red dots are emVars (n = 10) and gray dots are pCRE variants (n = 16). **f,** Proportion of MPRA inactive and pCRE variants, and emVars that are chromatin accessibility QTLs (caQTLs) from T cells[73]. **g,** Scatterplot comparing caQTL effect size (beta; x-axis) and MPRA log2 allelic bias (y-axis). Red dots are emVars ($n = 4$) and gray dots are pCREs ($n = 10$). **h,** Scatterplot comparing deltaSVM score (x-axis) with MPRA

$log_2$ allelic bias (y-axis). **i,** Proportion of MPRA inactive and pCRE variants and emVars that overlap TF motifs. **j,** Scatterplot comparing allele-specific TF binding scores (y-axis) and MPRA allelic bias (x-axis) for emVars predicted to perturb TF binding ($n = 9$). Calculations for (b and c) are risk ratios (see Methods) with Fisher's exact test *P*-values and Bonferroni correction (see Supplementary Tables 3 and 4 for exact *P*-values). (d, f, and i) *P* values calculated using two-sided two proportions z test with no multiple comparisons adjustment. (e, g, h and j) *P* values are from linear regression F statistic. (e, g, and j) shaded portions of the graph are the 95% confidence interval.

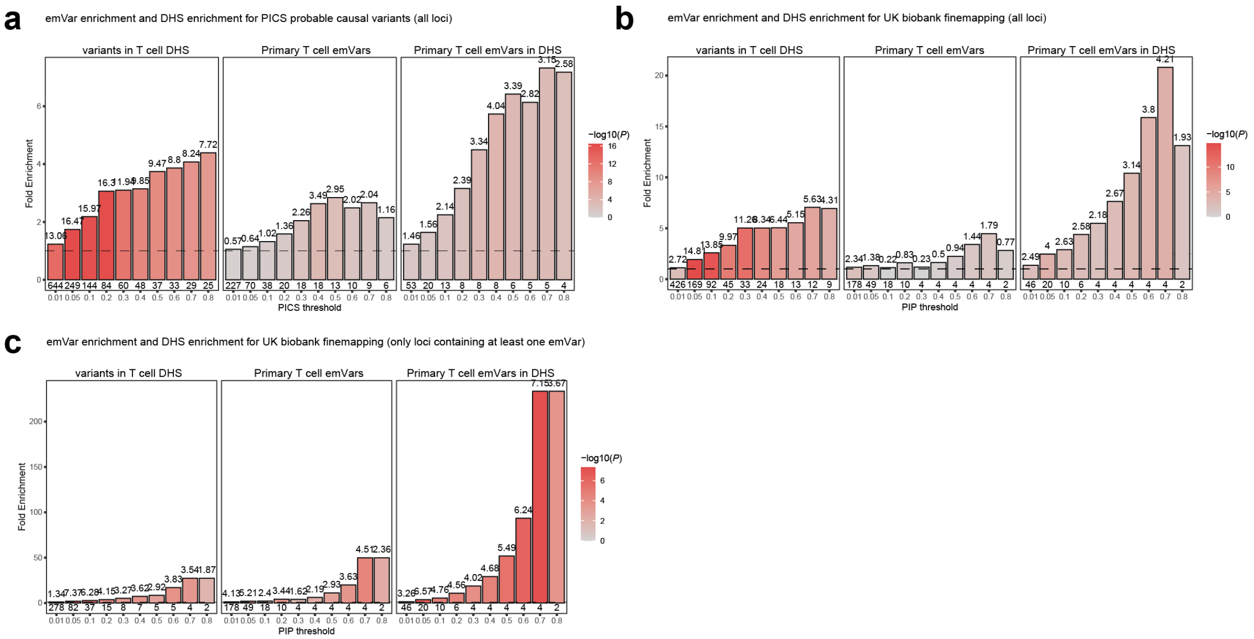


**Supplementary Fig. 5.**

**Primary T cell emVars enrich for causal variants. a-c,** Bar plot showing emVar enrichment for high posterior probability variants. Graphs in (a) and (b) consider all loci tested for PICS (a) and UKBB (b) fine mapping. The graph in (c) considers only loci with at least one emVar for UKBB finemapping. Each set of bar graphs is broken into three, with enrichment of fine-mapped variants within DHS sites alone (left), enrichment of fine-mapped variants that are primary T cell emVars (middle), and enrichment of fine-mapped variants that are emVars in T cell DHS sites (right), with the minimum posterior probability threshold indicated on the x-axis and fold enrichment shown on the y-axis. Details of PICS and UKBB enrichment results are shown in Supplementary Tables 8 and 9. Numbers below each bar show the number of emVars that are statistically fine-mapped at a given posterior probability threshold. Shade of each bar is the -$\log_{10}$ of the enrichment *P* value. Enrichment in (a-c) was calculated as a risk ratio (see Methods), and *P* values were determined through a two-sided Fisher's exact test.

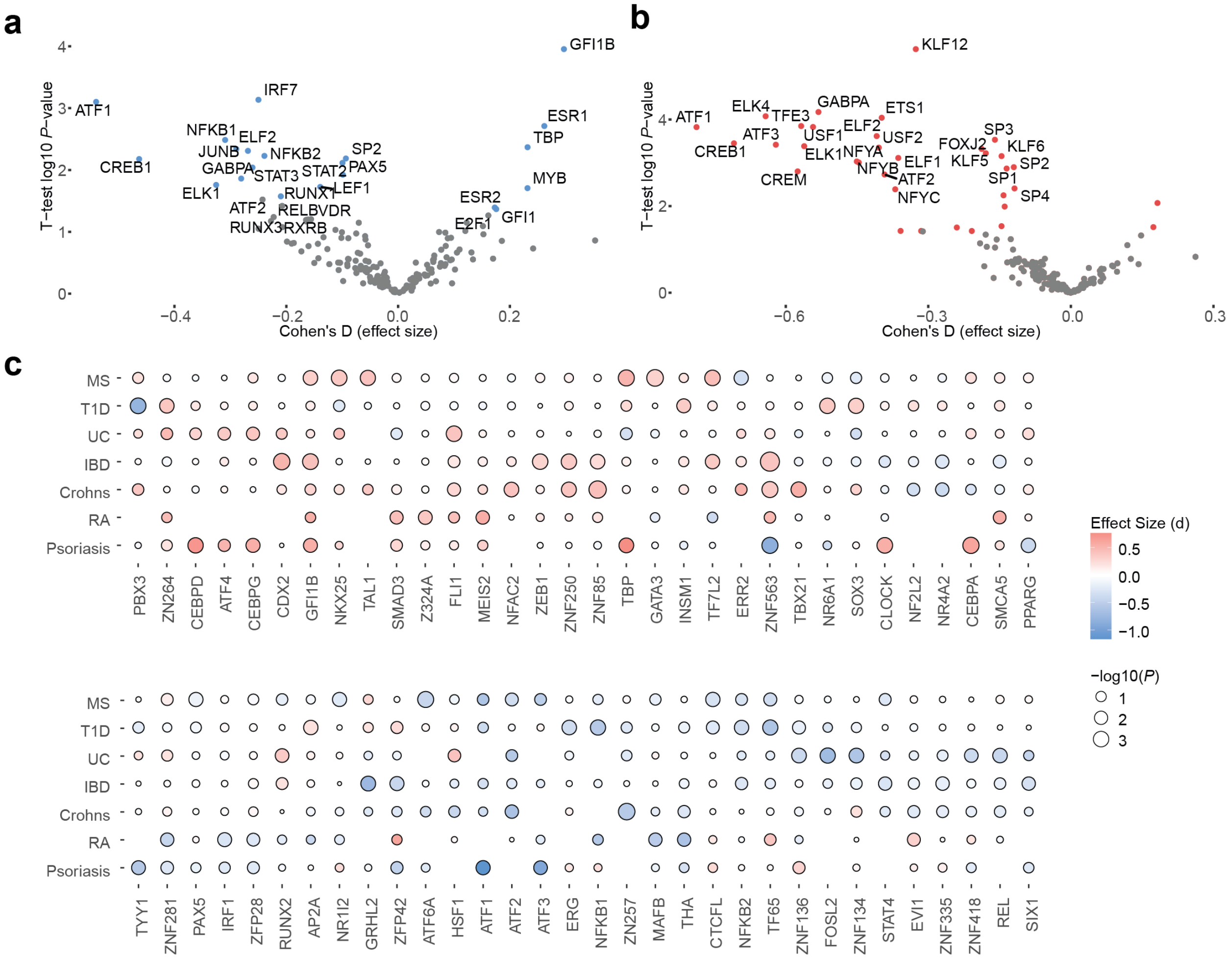

**Supplementary Fig. 6.**

**Primary T cell and Jurkat MPRAs identify different emVars modulated by distinct transcription factors. a and b,** Transcription factors whose motifs are predicted to be disrupted and the effect on allele-specific expression in (a) primary T cell MPRAs and (b) Jurkat MPRAs. Cohen's d on the x-axis shows the collective effect size of variant alleles that disrupt a given TF motif and -log$_{10}$ *P* value on the y-axis. **c,** TF motif disruption of variants by disease, with disease on the y-axis, TF motif on the x-axis, dot size the -log10 *P* values, and effect size (d) is color. The motifs whose disruption caused the most significant upregulation and downregulation of expression for each disease and were hierarchically clustered according to TF and disease. For A and B, effect size is calculated using Cohen's d for variant alleles predicted to disrupt a given TF motif and P values are calculated using a t-test comparing effect on expression of variants that disrupt a given motif versus all other variants. The -log *P* value in (a-c) is calculated using a *t* test comparing the effect on expression of variant alleles that disrupt a given motif compared to all other alleles tested.

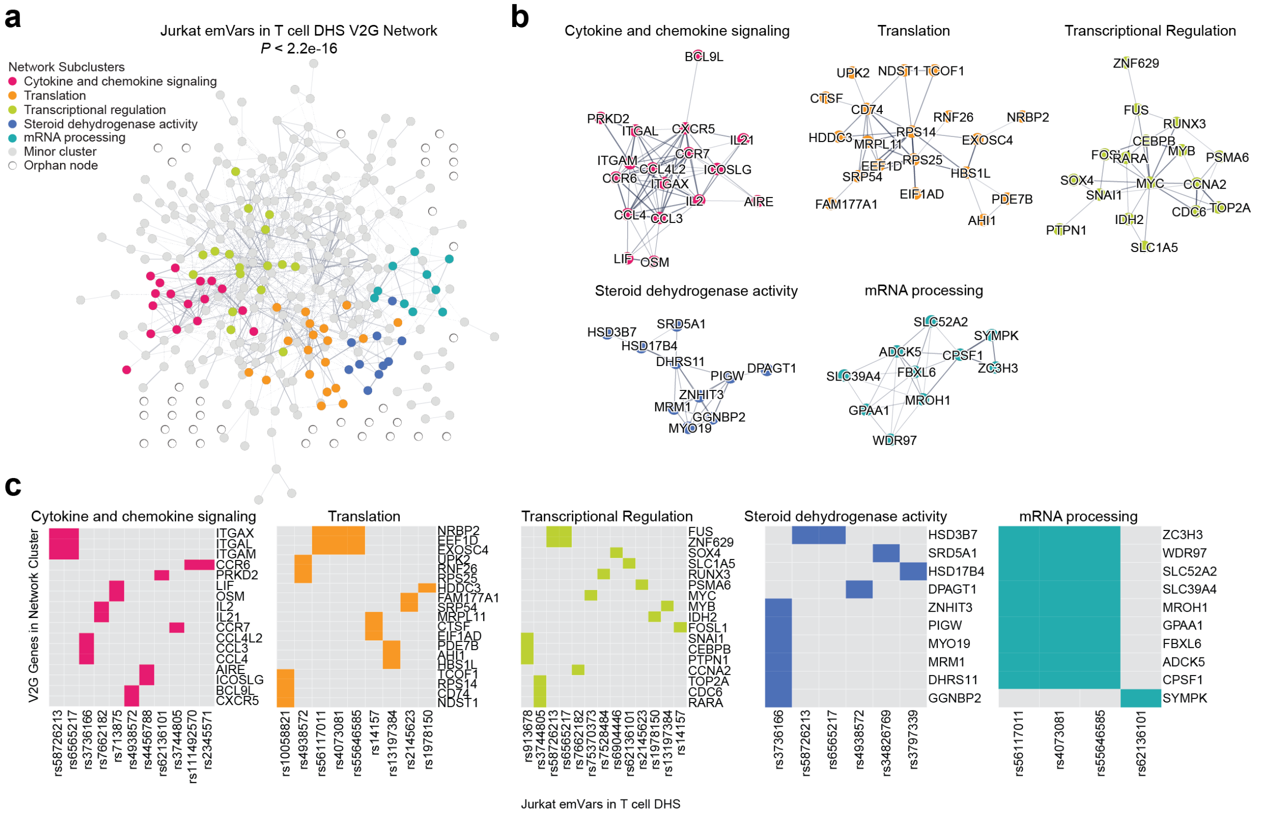

**Supplementary Fig. 7.**

**Network analysis of predicted target genes of Jurkat emVars. a,** STRING network showing V2G genes linked to 31 emVars in T cell DHS sites (nodes) and edges representing the strength of gene-gene interactions. **b,** The subclusters with the most genes from the larger network in (a) with labeled gene nodes. **c,** Top 5 Jurkat network clusters with each putative emVar on the x-axis and target gene on the y-axis. Fill color indicates that the gene is a V2G gene of the indicated emVar. *P*-value in (a) is calculated according to STRING protein-protein interaction enrichment given the expected versus observed number of edges within the network.

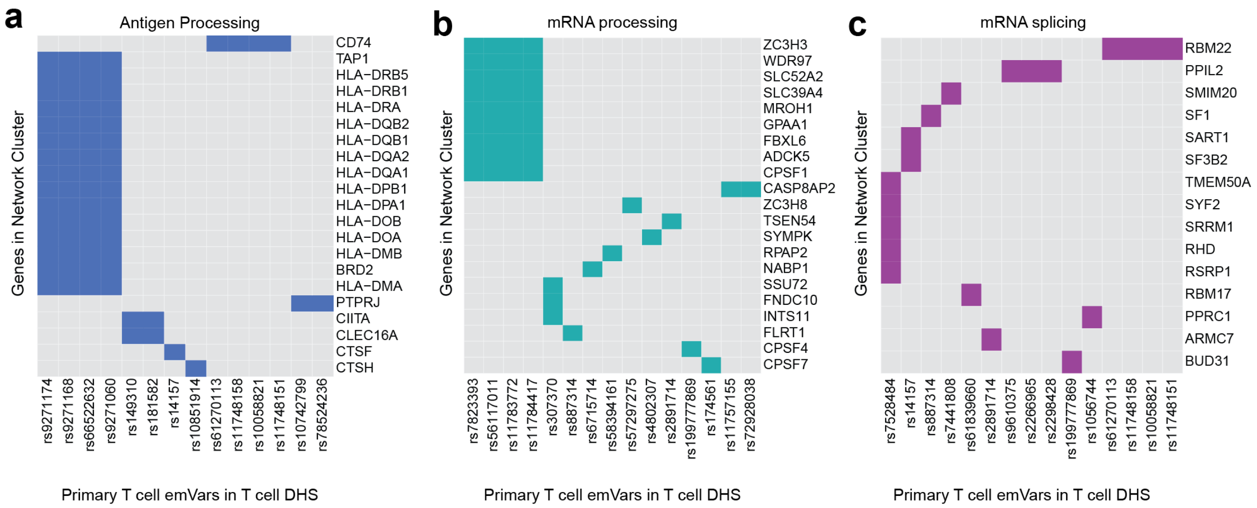


**Supplementary Fig. 8.**

**Other Primary T cell emVars in DHS network clusters**. **a-c,** The antigen processing (a), mRNA processing (b), and mRNA splicing (c) clusters from Fig. 3 with each putative emVar on the x-axis and target gene on the y-axis. Fill color indicates that the gene is a V2G gene of the indicated emVar.

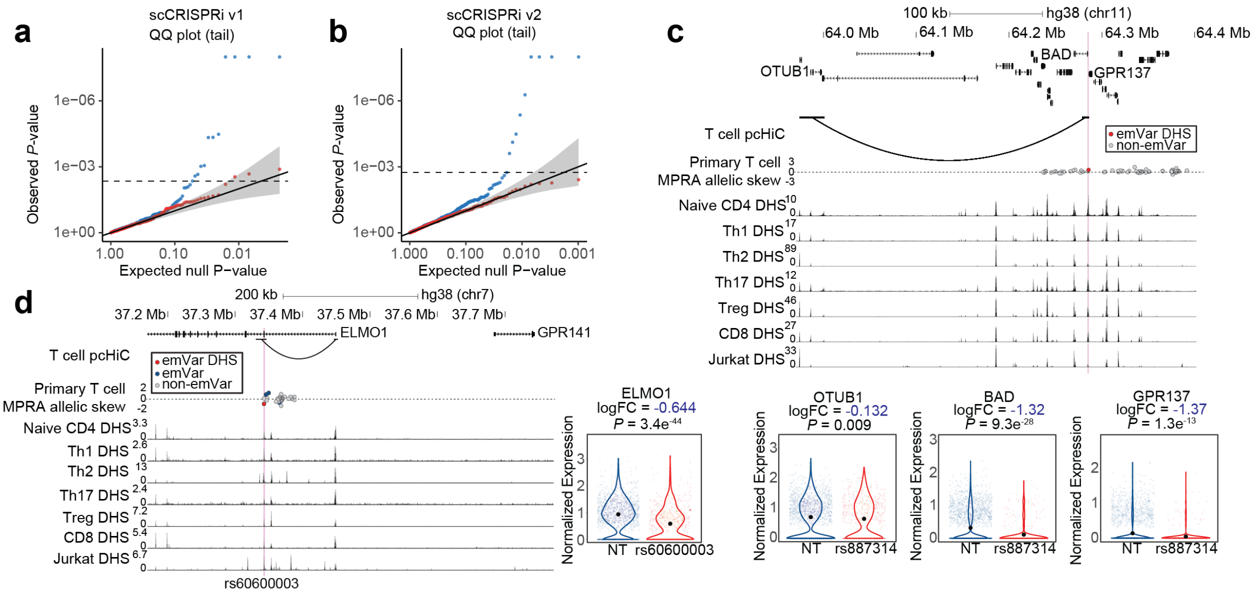


**Supplementary Fig. 9.**

**Single cell CRISPRi screens identify emVar target genes. a and b**, QQ plots showing the expected (x-axis) versus observed (y-axis) *P* value output of SCEPTRE, with dotted line indicating the significance cutoff for v1 (a) and v2 (b) libraries. **c and d,** Locus plots of the BAD (c) and ELMO1 (d) loci. pcHiC loops from primary human T cells are depicted below genes in the locus plot. Disease-associated variants (dots) are red if they are emVars in DHS, pink if they are emVars not within DHS, and gray if they are non-emVars. Accessible chromatin data from T cells are depicted as read pileups (peaks) on the locus track from various T cell types. The pink lines represent the location of emVars in DHS. Violin plots depicting genes that are differentially expressed when targeting CRISPRi to the emVar using gRNAs compared to cells containing non-target gRNAs. Shaded regions in (a and b) are the 95% confidence interval. Black dots in violin plots in (c and d) depict the mean of the distribution.

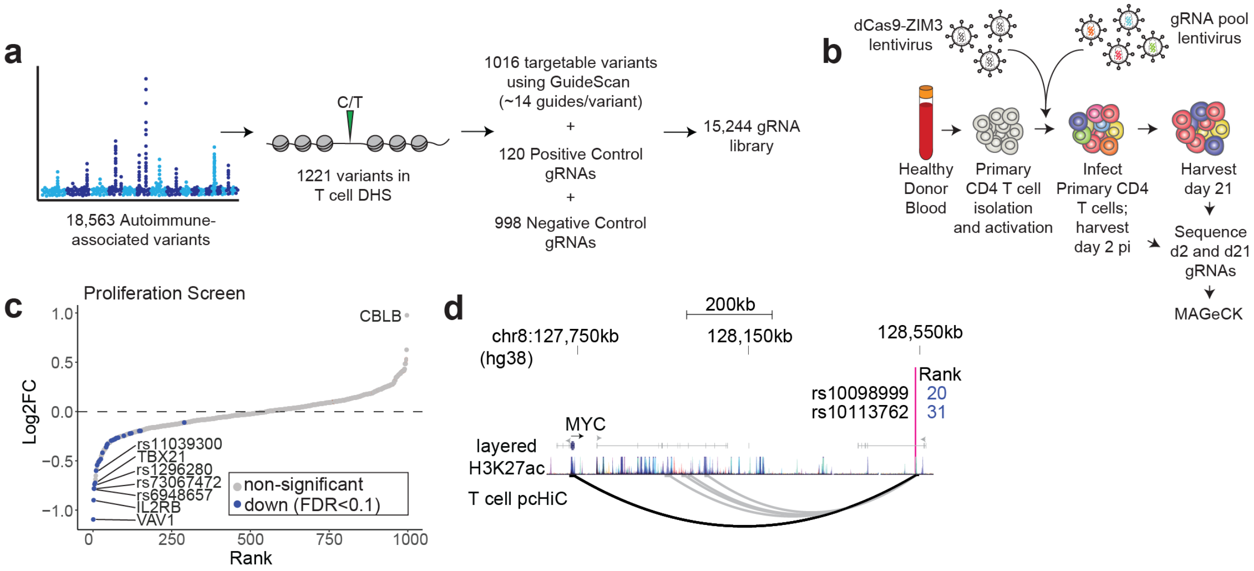


**Supplementary Fig. 10.**

**Genome-wide CRISPRi screen targeting variants in accessible chromatin. a,** Library makeup of genome-wide screen. **b,** Screen workflow. **c,** Rank order plot depicting targets of the CRISPRi screen, with positive control genes VAV1, IL2RB, TBX21, and CBLB and variants indicated by rsid. *P*-values are determined using MAGeCK. **d,** Locus plot of the MYC locus showing two variants that are proliferation hits in the screen (rank of hit in blue) in a distal enhancer contact the MYC promoter as determined by pcHiC in T cells.

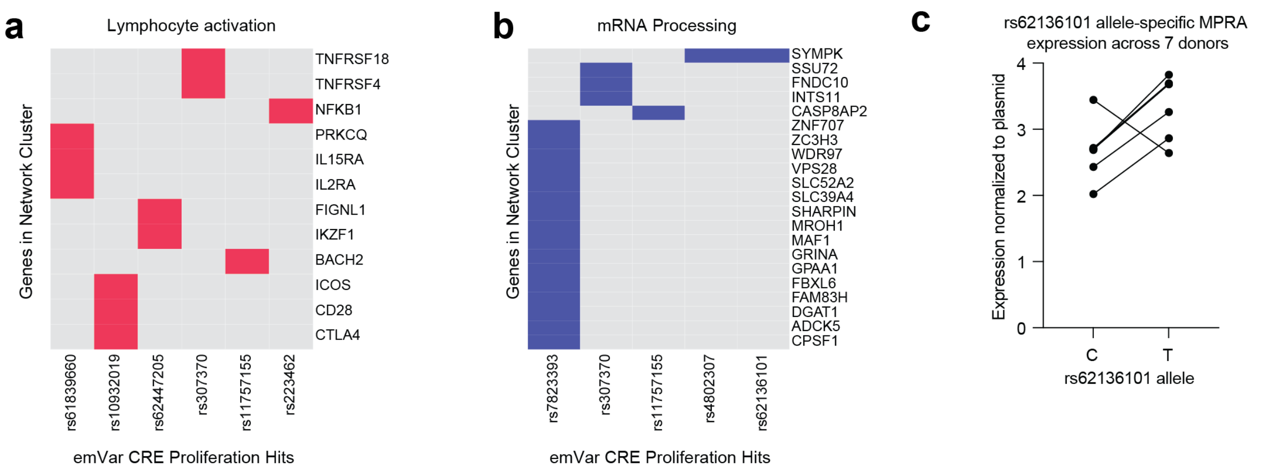


**Supplementary Fig. 11.**

**emVars targeting genes within proliferation screen network clusters. a and b,** The lymphocyte activation (a) and mRNA processing (b) clusters with each putative emVar target gene represented with color. **c,** Normalized counts of rs62136101 variant alleles tested in primary T cell MPRAs across 7 donors. Each line indicates one donor.

**Supplementary Table 1. Primary T cell MPRA results.**

Primary T cell MPRA results. All MPRA expression and allelic bias (skew) results. *P* values were calculated using generalized linear models and multiple testing was corrected using FDR calculated using the method of Benjami and Hochberg.

**Supplementary Table 2. Jurkat MPRA results.**

Jurkat cell line MPRA results. All MPRA expression and allelic bias (skew) results. *P* values were calculated using generalized linear models and multiple testing was corrected using FDR calculated using the method of Benjami and Hochberg.

**Supplementary Table 3. chromHMM enrichment of emVars and pCREs for primary T cell MPRA.**

chromHMM enrichments. Fold enrichments and *P* values for enrichment of chromHMM partitions in pCREs or emVars from the primary T cell MPRA. *P* values shown are calculated using two-sided Fisher's exact test.

**Supplementary Table 4. Epigenetic modification enrichment of emVars and pCREs for primary T cell MPRAs.**

Histone mark, CAGE, and DHS enrichments. Fold enrichments and *P* values for enrichment for various T cell histone marks, CAGE annotations, and DHS sites in pCREs or emVars in the primary T-cell MPRA. *P* values shown are calculated using two-sided Fisher's exact test.

**Supplementary Table 5. Primary T cell MPRA functional annotations.**

MPRA functional annotation. T cell MPRA effect, DHS, ATAC allelic skew, caQTL, T cell eQTL, Ananastra and TF motifbreakR information for each variant.

**Supplementary Table 6. PICS fine-mapping results for MPRA-tested variants.**

PICS fine-mapping of MPRA variants. Results of PICS fine-mapping for MS, RA, T1D, psoriasis, IBD, ulcerative colitis (UC), and Crohn's on MPRA variants.

**Supplementary Table 7. UKBB fine-mapping data for MPRA-tested variants.**

UKBB fine-mapping of MPRA variants. Results of UKBB fine-mapping for all UKBB traits on MPRA variants.

**Supplementary Table 8. PICS fine-mapped variant enrichment for primary T cell MPRA data.**

emVar PICS enrichment for all loci in the primary T cell MPRA and loci containing an emVar. *P* values shown are from a two-sided Fisher's exact test.

**Supplementary Table 9. UKBB fine-mapped variant enrichment for primary T cell MPRA data.**

emVar UKBB enrichment for all loci in the primary T cell MPRA and loci containing an emVar. *P* values shown are from a two-sided Fisher's exact test.

**Supplementary Table 10. motifbreakR results for primary T cell MPRA data.**

Primary T cell MPRA and motifbreakR TF motif disruption. Details on all primary T cell MPRA variants with significant predicted disruption of TF position-weighted matrices based on motifbreakR run with HOCOMOCO v.11.

**Supplementary Table 11. Statistical test for effect of variants that break motifs on primary T cell MPRA allele-specific expression.**

motifbreakR motif disruption vs. primary T cell MPRA skew. Results of the Student's *t* test and Cohen's d comparing the Primary T-cell MPRA skew (allele specific differences in expression) of variants which motifibreakR indicates disrupts versus does not disrupt a specific TF motif.

**Supplementary Table 12. motifbreakRresults for Jurkat MPRA data.**

Jurkat cell line MPRA and motifbreakR TF motif disruption. Details on all Jurkat cell line MPRA variants with significant predicted disruption of TF position-weighted matrices based on motifbreakR run with HOCOMOCO v.11.

**Supplementary Table 13. Statistical test for effect of variants that break motifs on Jurkat T cell MPRA allele-specific expression.**

motifbreakR motif disruption vs. Jurkat cell line MPRA Skew. Results of the Student's *t* test and Cohen's d comparing the Jurkat cell line MPRA skew (allele specific differences in expression) of variants which motifibreakR indicates disrupts versus does not disrupt a specific TF motif.

**Supplementary Table 14. Variant to Gene tables for primary T cell emVars in accessible chromatin.**

Open Targets V2G genes for 79 Primary T cell emVars in T cell accessible chromatin. Genes are filtered for expression in primary T cells (TPM > 1). V2G data are based on distance to variant, eQTL, sQTL, pQTL, and pcHiC data.

**Supplementary Table 15. Variant to Gene tables for Jurkat emVars in accessible chromatin.**

Open Targets V2G genes for 31 Jurkat emVars in T cell accessible chromatin. Genes are filtered for expression in primary T cells (TPM > 1). V2G data are based on distance to variant, eQTL, sQTL, pQTL, and pcHiC data.

**Supplementary Table 16. Primary T cell emVars in T cell DHS sites STRING network clusters.**

Genes listed in STRING clusters of 79 primary T cell emVars in T cell accessible chromatin.

**Supplementary Table 17. EnrichR Panther module for primary T cell emVars in T cell DHS site V2Gs.**

EnrichR Panther module enrichment of network in Fig. 3.

**Supplementary Table 18. Jurkat emVars in T cell DHS sites STRING network clusters.**

Genes listed in STRING clusters of 31 Jurkat emVars in T cell-accessible chromatin.

**Supplementary Table 19. Single cell CRISPRi library v1.**

Single cell CRISPRi library v1 targeting 20 emVars and 3 pCREs in accessible chromatin with PICS>0.1.

**Supplementary Table 20. Single cell CRISPRi library v2.**

Single cell CRISPRi gRNA library v2 targeting 50 emVars in accessible chromatin, preferencing those that are > 3500 bp from the TSS.

**Supplementary Table 21. Single cell CRISPRi results v1.**

SCEPTRE results of scCRISPRi v1.

**Supplementary Table 22. Single cell CRISPRi results v2.**

SCEPTRE results of scCRISPRi v2.

**Supplementary Table 23. Single cell CRISPRi STRING network.**

STRING clusters of genes that are differentially expressed when targeting CRISPRi to emVars in accessible chromatin in scCRISPRi screens.

**Supplementary Table 24. Genome-wide (~1000 variant) bulk CRISPRi proliferation screen library.**

Proliferation screen gRNA library targeting ~1000 autoimmune-associated variants in accessible chromatin.

**Supplementary Table 25. Analysis on proliferation screen of ~1000 variant putative CREs.**

MAGeCK analysis of full genome-wide (~1000 variant CRE) proliferation screen.

**Supplementary Table 26. Analysis on proliferation screen focused on 56 emVars in T cell DHS sites.**

MAGeCK analysis focusing on 56 emVars in T cell-accessible chromatin.

**Supplementary Table 27. Proliferation screen STRING network.**

STRING clusters of V2G genes of emVars in T cell-accessible chromatin that are hits within the proliferation screen.

**Supplementary Table 28. Open Targets V2G genes for 13 primary T cell emVars in T cell DHS sites that are proliferation hits.** Genes are filtered for expression in primary T cells (TPM > 1). V2G data are based on distance to variant, eQTL, sQTL, pQTL, and pcHiC data.

**Supplementary Table 29. Differential gene expression results comparing gRNAs targeting the PPP5C TSS versus a non-targeting gRNA.**

DESeq2 output of CRISPRi+ T cells that contain PPP5C TSS-targeted gRNAs versus an NT gRNA.

**Supplementary Table 30. Differential gene expression results comparing gRNAs targeting the rs62136101 CRE versus a non-targeting gRNA.**

DESeq2 output of CRISPRi+ T cells that contain rs62136101-targeted gRNAs versus an NT gRNA.

**Supplementary Table 31. Primers, antibodies, and other reagents used in this study.**

Primers and antibodies used in this study.

**Supplementary Table 32. T cell DHS gridsearch statistics.**

Statistical results from grid search conducted on primary T cell MPRA data.

**Supplementary Table 33. Jurkat DHS gridsearch statistics.**

Statistical results from grid search conducted on Jurkat MPRA data.

**Supplementary Table 34. ENCODE DHS enrichment of pCRE elements in primary T cell MPRAs**

Statistical results from enrichment calculations of pCRE elements for ENCODE DHS sites from all tested cell types.

**Supplementary Table 35. Statistical test for effect of variants that break motifs on primary T cell MPRA allele-specific expression by each disease.**

motifbreakR motif disruption vs. primary T cell MPRA skew broken down by disease. Results of the Student's *t* test and Cohen's d comparing the Primary T-cell MPRA skew (allele specific differences in expression) of variants which motifibreakR indicates disrupts versus does not disrupt a specific TF motif.

**Supplementary Data 1. Open Targets V2G genes for all variants tested in the MPRA library.**

Open Targets V2G genes for all variants in this study. Genes are filtered for expression in primary T cells (TPM>1). V2G data are based on distance to variant, eQTL, sQTL, pQTL, and pcHiC data.
